# PREFRONTAL CORRELATES OF FEAR GENERALIZATION DURING ENDOCANNABINOID DEPLETION

**DOI:** 10.1101/2024.01.30.577847

**Authors:** Luis E. Rosas-Vidal, Saptarnab Naskar, Leah M. Mayo, Irene Perini, Megan Altemus, Hilda Engelbrektsson, Puja Jagasia, Markus Heilig, Sachin Patel

**Affiliations:** Northwestern University, Feinberg School of Medicine, Department of Psychiatry and Behavioral Sciences, Chicago, IL; Hotchkiss Brain Institute and Mathison Centre for Mental Health Research and Education, Department of Psychiatry, Cumming School of Medicine, University of Calgary, Calgary, Alberta, Canada; Linköping University, Center for Social and Affective Neuroscience, Department of Biomedical and Clinical Sciences, Linköping, Sweden; Vanderbilt University Medical Center, Department of Psychiatry and Behavioral Sciences, Nashville, TN

## Abstract

Maladaptive fear generalization is one of the hallmarks of trauma-related disorders. The endocannabinoid 2-arachidonoylglycerol (2-AG) is crucial for modulating anxiety, fear, and stress adaptation but its role in balancing fear discrimination versus generalization is not known. To address this, we used a combination of plasma endocannabinoid measurement and neuroimaging from a childhood maltreatment exposed and non-exposed mixed population combined with human and rodent fear conditioning models. Here we show that 2-AG levels are inversely associated with fear generalization at the behavioral level in both mice and humans. In mice, 2-AG depletion increases the proportion of neurons, and the similarity between neuronal representations, of threat-predictive and neutral stimuli within prelimbic prefrontal cortex ensembles. In humans, increased dorsolateral prefrontal cortical-amygdala resting state connectivity is inversely correlated with fear generalization. These data provide convergent cross-species evidence that 2-AG is a key regulator of fear generalization and suggest 2-AG deficiency could represent a trauma-related disorder susceptibility endophenotype.

## INTRODUCTION

Generalization of conditioned associations is advantageous for survival as it maximizes goal attainment while minimizing harm, based on rewarding or dangerous experiences, respectively. However, overgeneralization is a hallmark and often debilitating symptom of psychiatric disorders including post-traumatic stress disorder (PTSD) ^1–5^, and underlies broadening of sensory stimuli capable of inducing hyperarousal and avoidance symptoms central to the disorder. The amygdala and the medial prefrontal cortex (mPFC) are crucial for balancing fear memory generalization and discrimination ^3,6–8^. Inactivation or lesions of the mPFC increase contextual fear generalization ^9,10^ and generalization states correlate with loss of mPFC-amygdala entrainment ^11^. Specifically, in rodents, the prelimbic prefrontal cortex (PL) plays a crucial role in fear expression ^12,13^ and opposing generalization of the conditioned stimuli to the neutral stimuli ^8,14,15^. It is not known, however, how generalization between discrete fear-associated and neutral stimuli is represented by neuronal ensembles within the PL.

The endocannabinoid system has been implicated in the regulation of stress, anxiety, and fear states and has been proposed as a promising target for stress-related psychiatric disorders ^16^. One of the endocannabinoids, 2-arachidonoylglycerol (2-AG), is synthesized primarily by diacylglycerol-lipase alpha (DAGLα) in the brain and inhibition of this enzyme in experimental animals is associated with increased anxiety, decreased resiliency to stress, and impaired fear extinction ^17–20^. While 2-AG has been implicated in stress resiliency and found to be lower in the plasma of subjects with PTSD ^21^ but see ^22^, the association between 2-AG levels and fear generalization has not been systematically investigated in humans or animal models.

In this study, we use fear conditioning in both mice and humans in combination with plasma 2-AG measurements, pharmacological manipulations, human brain imaging, and single neuron calcium imaging to establish the role of 2-AG levels in the regulation of fear generalization and its impact on prefrontal neural correlates. We show that 2-AG levels predict and regulate fear memory generalization and that sensory stimuli specificity within PL neuronal ensembles deteriorates in 2-AG deficient states. Furthermore, resting state prefrontal-amygdala connectivity is inversely correlated with fear generalization. These data provide new insights into 2-AG regulation of fear generalization in humans and mice and reveal how 2-AG deficient states affect cellular correlates of fear generalization in PL neurons.

## METHODS

### Study Overview

This study consisted of one screening visit and two laboratory sessions. During screening, participants were evaluated for eligibility and complete questionnaire measures, including the Difficulties in Emotion Regulation Scale ^23^ (see Figure 1a for study schematic). In the first laboratory session, blood samples and psychophysiological recordings were collected and participants completed behavioral tasks, including fear conditioning and extinction. The second session included a magnetic resonance imaging (MRI) session, with one anatomical, one post-recall resting-state, and three task-based scans (other tasks not included here, see ^24^). The study was approved by the Regional Ethics Review Board in Linköping, Sweden (Dnr 2015/256-31, and 2017/41-32). All behavioral data were analyzed using the Statistical Package for Social Sciences (SPSS) software version 28.0.1.0 and graphs were created in Prism 9.

**Fig. 1.**
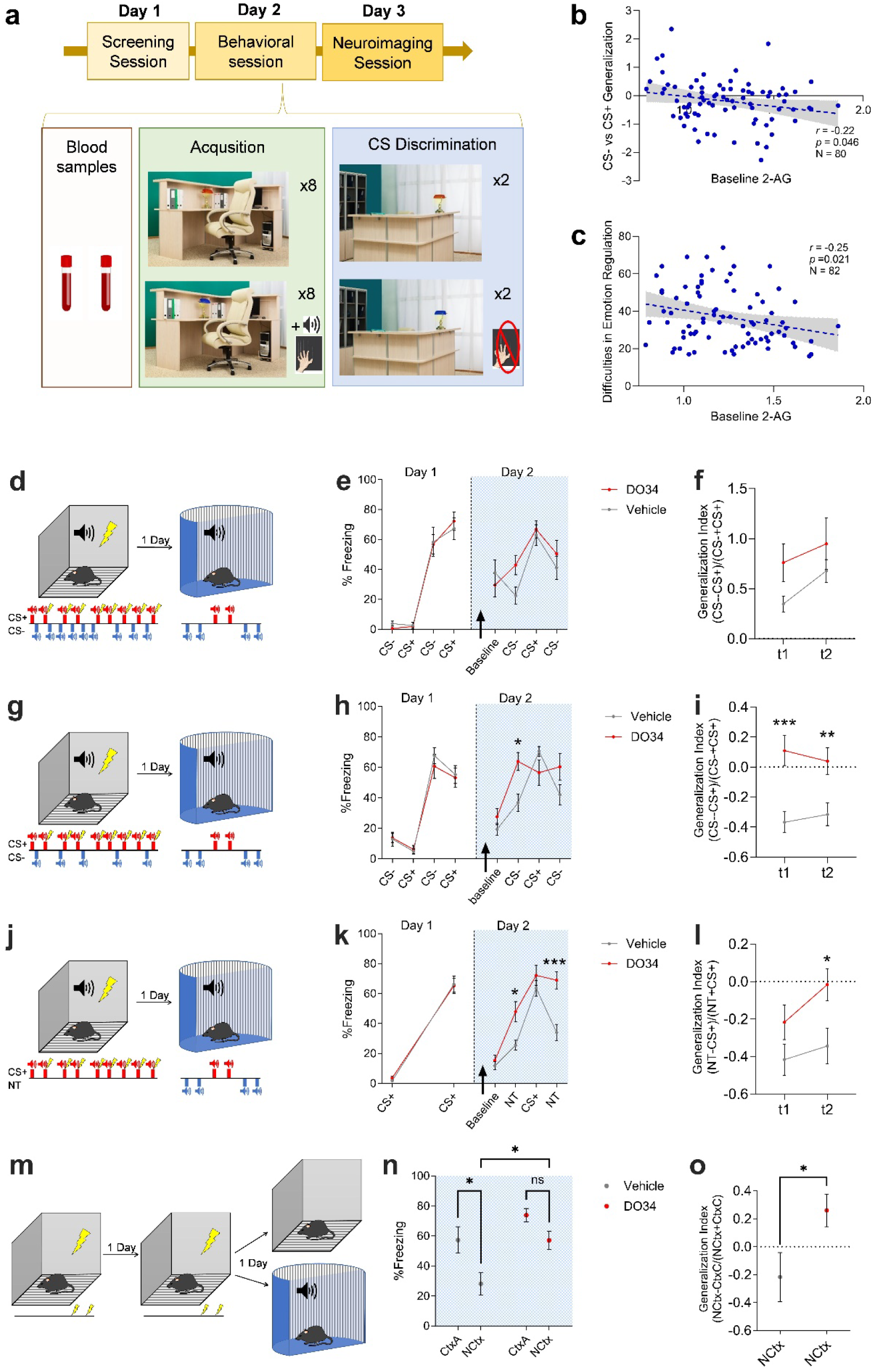
Decreased levels of 2-AG are associated with increased generalization to neutral stimuli and self-reported emotional regulation. (a) Representative schematic of experiment. All participants completed three sessions on three separate days. (b) Discrimination between conditioned (CS+) and safe (CS-) stimuli as a function of baseline peripheral 2-AG levels. (c) Self-reported difficulties in emotional regulation (Difficulties in Emotion Regulation Scale) as a function of baseline peripheral 2-AG levels. (d) Schematic of experiment with depiction of stimuli presented during differential fear conditioning and recall test sessions. (e) % time freezing to CS+ and CS-tones. Arrow depicts i.p. injection of vehicle or DO34 prior to recall test on Day 2. (f) Generalization index for CS-tones t1 and t2 for vehicle and DO34 exposed groups. (g) Schematic of experiment with depiction of stimuli presented during partial differential fear conditioning and recall test sessions. (h) % time freezing to CS+ and CS-tones. (i) Generalization index for CS-tones t1 and t2 for vehicle and DO34 exposed groups. (j) Schematic of experiment with depiction of stimuli presented during classical fear conditioning and recall test sessions. (k) % time freezing to CS+ and NT. Arrow depicts i.p. injection of vehicle or DO34 prior to recall test on Day 2. (l) Generalization index for NT t1 and t2 for vehicle and DO34 exposed groups. (m) Schematic of experiment with depiction of stimuli presented during contextual fear conditioning and recall test sessions. Arrow depicts i.p. injection of vehicle or DO34 prior to recall test on Day 3. (n) Freezing to conditioning context (CtxA) and novel context (NCtx). (o) Generalization index calculated using freezing of test context relative to freezing to Ctx A on conditioning Day 2 (CtxC). Repeated measures ANOVA or two-way ANOVA followed by Šídák’s multiple comparisons test where appropriate. *p < 0.05, **p<0.01, ***p<0.001.

### Human Participants

Participants were recruited between March 2017 and July 2020, at Linköping University. A total of 101 participants were included in the study, divided into four groups across the dimensions of histories of childhood maltreatment (CM) and substance use disorders (SUD), described further in Supplementary Methods and ^24,25^. However, there were no group differences across the measures described below, so all analyses were collapsed across groups for a total N = 101 (see Table 1 for demographic information). Of note, individuals with histories of CM were recruited via prior contact with a specialized trauma unit within Child & Adolescent Psychiatry at the regional hospital in Linköping, which was created in the early 1990’s. As a result, the upper age limit of patients was dependent on being a former patient of the trauma unit since its inception, while the lower age limit was 18.

**Table 1:**
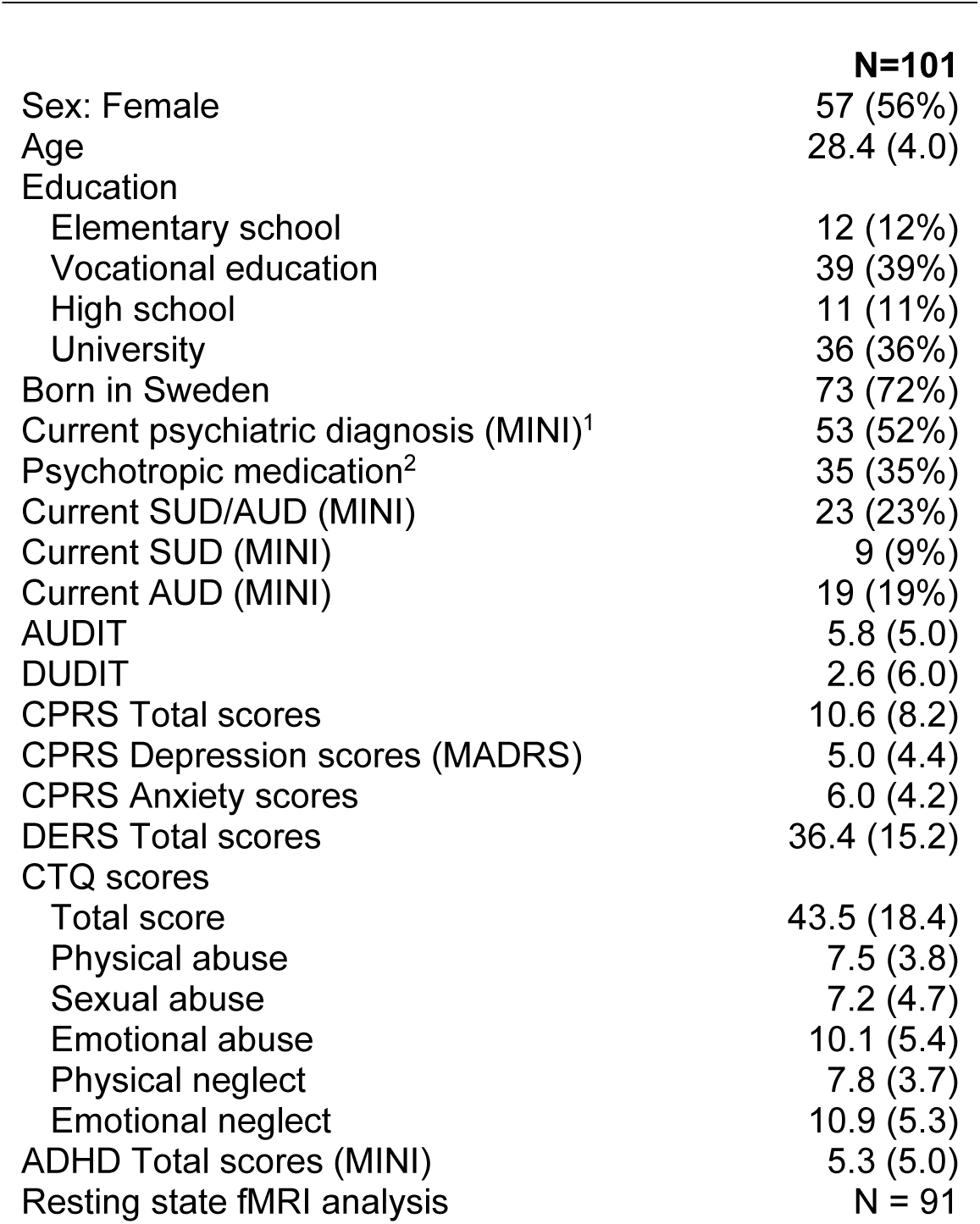
Demographic Characteristics. Values represent mean (± standard deviation) or N (percent of N). MINI = Mini International Neuropsychiatric Interview; SUD = substance use disorder; AUDIT = Alcohol Use Disorder Identification Test; DUDIT = Drug Use Disorder Identification Test; CPRS = Comprehensive psychopathological rating Scale; MADRS = Montgomary-Åsberg Depression Rating Scale; CTQ = Childhood Trauma Questionnaire; ADHD = attention deficit hyperactivity disorder; fMRI = functional magnetic resonance imaging

Participants completed breath and urine screens for alcohol and drug use prior to laboratory sessions. During the screening session, participants underwent a psychiatric clinical assessment by a trained research nurse or study physician. After determining eligibility, participants received research information, provided written informed consent, and completed self-report questionnaires. Detailed description can be found in Supplementary Methods.

### Human Fear Conditioning

Upon arrival at the lab, participants were fitted with an intravenous catheter for blood sample collection for subsequent analysis of endocannabinoid levels and prepared for psychophysiological recordings via application of facial electromyography (EMG) recording electrodes on the orbicularis muscle to assess the eyeblink component of the startle response. EMG data collection and fear conditioning took place as previously described ^26,27^ and provided in detail in supplementary methods. Briefly, the fear conditioning task consisted of Habituation, Acquisition, and Recall, with the first 4 trials of Extinction serving as the CS generalization test ^28^. In Acquisition, a specific conditioned stimulus (lamp color, CS+) predicted an unpleasant sound (nails across the chalkboard; US ^29^ while another CS (different lamp color, CS-) was not followed by any stimulus. After a 10min break, Recall took place in a different context (different picture on the computer) with the same CS+ and CS-but no US. A startle probe was used to elicit blinks during each CS and inter-trial intervals, measured as the peak-to-peak EMG value. To account, startle responses to the CS+ were standardized to the mean startle response during ITI/rest trials to account for individual differences in startle magnitude ^26,27,30,31^. Due to technical issues (i.e., inadvertent partial removal of the EMG sensor during task completion), N = 3 individuals did not have complete fear conditioning data and thus were not included in the analysis.

### Endocannabinoid analysis

Baseline endocannabinoids levels were calculated as the average of two timepoints. AEA and 2-AG were extracted and analyzed using liquid chromatography tandem mass spectrometry (LC-MS/MS), as previously published ^26,32^ (Supplementary Methods). Endocannabinoid values were log transformed due to non-normality of the distribution; these transformed values were used in all subsequent analyses. Baseline differences in endocannabinoids were added as co-variates in analysis of brain, behavioral, and self-report measures. Of the N = 101 participants, technical issues prevented blood from being drawn in N = 13. An additional N = 6 had 2-AG levels below the lower limit of detection. Thus, analyses of eCB levels included N = 82 participants. A total of N = 3 participants were removed from the fear conditioning analysis (of them, N = 1 did not have eCB data). Thus, N = 80 participants had complete fear conditioning and eCB data.

### Magnetic Resonance Imaging

Anatomical and functional blood oxygen-level-dependent (BOLD) data were collected on a Siemens MAGNETOM Prisma 3T MRI scanner (Siemens healthcare AB, Stockholm, Sweden) equipped with a 64-channel head coil. EPI images were de-spiked, slice-time corrected, smoothed (4 mm) and motion-corrected. Preprocessing and statistical analysis of resting state data were performed with the Analysis of Functional Neuro Images (AFNI) software v18.3.16 ^33^.

For each participant (N = 88), amygdala time course was entered as predictor in a regression analysis on preprocessed data, using 3dDeconvolve. Resulting 3D volumes with beta coefficients for right/left amygdala were assessed for covariance with peripheral measures of endocannabinoid function and behavioral measures of fear conditioning using the AFNI function 3dttest++ and with 2AG (N = 82) and CS generalization (CS-vs CS+) at early Extinction (N = 88) as covariates in two separate analyses. Activation maps were thresholded at per-voxel p < 0.002, cluster corrected at α = 0.05 ^34^. To validate the results, beta coefficients from the significant cluster were then extracted for each participant and entered in regression analyses with the corresponding covariate as predictor. Regression analyses were performed using SPSS version 25.

### Rodents

All experiments were approved by the Vanderbilt University and Northwestern University Institutional Animal Care and Use Committees and were conducted in accordance with the National Institute of Health guidelines for the Care and Use of Laboratory Animals. A total of 127 male C57BL/6J mice older than 8 weeks and obtained from Jackson Labs or bred in our animal facility, were used in our experiments. Mice were housed in a temperature and humidity-controlled housing facility under a 12h light/dark cycle with ad libitum access to food. Mice were group housed except for mice implanted with Gradient Index (GRIN) lenses which were single housed. Statistical analysis and graphing were performed using GraphPad Prism 9 and the analysis performed is described in figure legends as appropriate.

### Rodent Surgeries

Briefly, mice were anesthetized with isofluorane and mounted into a stereotax. Skull hairs were trimmed, and overlying skin was cleaned with 70% isopropyl alcohol and iodine. A midline incision was performed, and a small hole was drilled above the PL to allow for injection of AAV encoding GCaMP7f under control of the synapsin promoter (AAVrg-syn-jGCaMP7f-WPRE). Following injection, a GRIN lens was lowered into PL and was cemented to the skull. Mice were allowed to recover for at least 2 weeks. More detailed information can be found in Supplementary Methods and Patel et al., 2022.

### Rodent Behavior and recordings

Mice were exposed to fear conditioning protocols as depicted in Figure 2 and described in detail in Supplementary Methods. For auditory fear conditioning, foot shocks were 0.4 mA in intensity and 1s in duration. 2 kHz and white noise tones at 75 dB were counterbalanced across experiments to control for stimulus effects. For contextual fear conditioning, mice were exposed to 2 sessions in the conditioning context (CtxA), 270 s in duration, 1 per day, with 2 0.4 mA foot shocks, 2 s in duration. Mice were injected with DO34 prior to recall phase of conditioning experiments. Recall phase was conducted 2 hours following DO34 injections and was carried out in a modified conditioning chamber (CtxB). For contextual fear recall, mice were exposed to the modified context as the novel context (NCtx) or to CtxA.

**Fig. 2.**
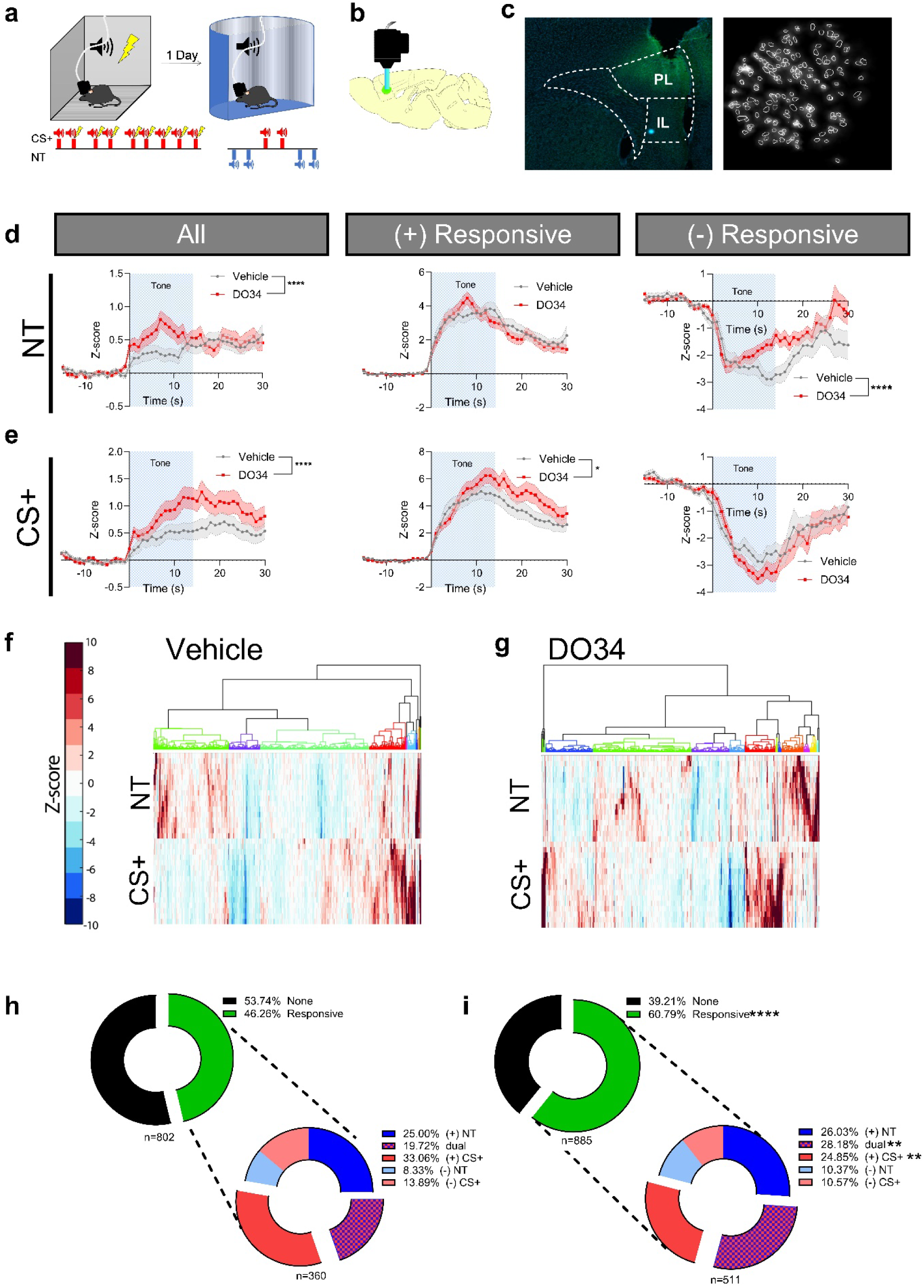
Neuronal tone responses to novel tones and CS+. (a) Schematic of experimental timeline and setup. (b) Sagittal depiction of GRIN lens and microscope implanted above GCaMP expression in the prelimbic cortex. (c) Left. Example coronal section of lens track above GCaMP expression in PL. Right. Maximum projection image as seen through the miniaturized microscope with identified neurons extracted with CNMFe algorithm surrounded by white lines. (d) Left. Average PL neuronal response to novel tone (NT) for all recorded neurons. Middle. NT response for neurons in PL that exceed +3 Zs during tone presentation. Right. CS+ response for neurons in PL below -3 Zs during tone presentation. (e) Left. Average PL neuronal response to conditioned tone (CS+) for all recorded neurons. Middle. CS+ response for neurons in PL that exceed +3 Zs during tone presentation. Right. CS+ response for neurons in PL below -3 Zs during tone presentation. (f) Hierarchical tree clustering of Z-scored neuronal activity to NT and CS+ from vehicle and DO34 exposed animals. (h) Pie charts for vehicle and (i) DO34 exposed animals showing proportion of neurons that significantly respond to tone presentations (±3 Zs). Bottom. Proportion of neurons that increase activity to NT ((+)NT) or CS+ ((+)CS+), decrease activity to NT ((-)NT) or CS+ ((-)CS+), or change activity to both NT and CS+ (dual). Data shown in mean ± SEM. mean ± SEM. Repeated measures ANOVA followed by Šídák’s multiple comparisons test and Fisher’s exact test where appropriate. *p < 0.05, **p<0.01, ***p<0.001, ****p<0.0001. n= 802 and 885 neurons from 7 and 8 mice treated with vehicle or DO34 respectively.

Calcium imaging experiments were conducted during fear conditioning as described above and depicted Figure 3a. The miniaturized microscope (Inscopix, Palo Alto, CA) was attached prior to behavioral sessions. On recall day, home cage recordings were conducted for 15 minutes before and 2 hours after DO34 injections. Following home cage recording sessions, mice were transferred to CtxB and exposed to the recall protocol. Data was acquired at a sampling rate of 10 Hz. All the fear conditioning experiments were controlled using FreezeFrame (Actimetrics, Wilmette, IL). For calcium activity recordings, the calcium imaging data acquisition system was interfaced with FreezeFrame using custom made RJ12 connector to BNC connector cables to deliver TTL pulses synchronizing the start of the experiment with the start of data acquisition and tone deliveries.

**Fig. 3.**
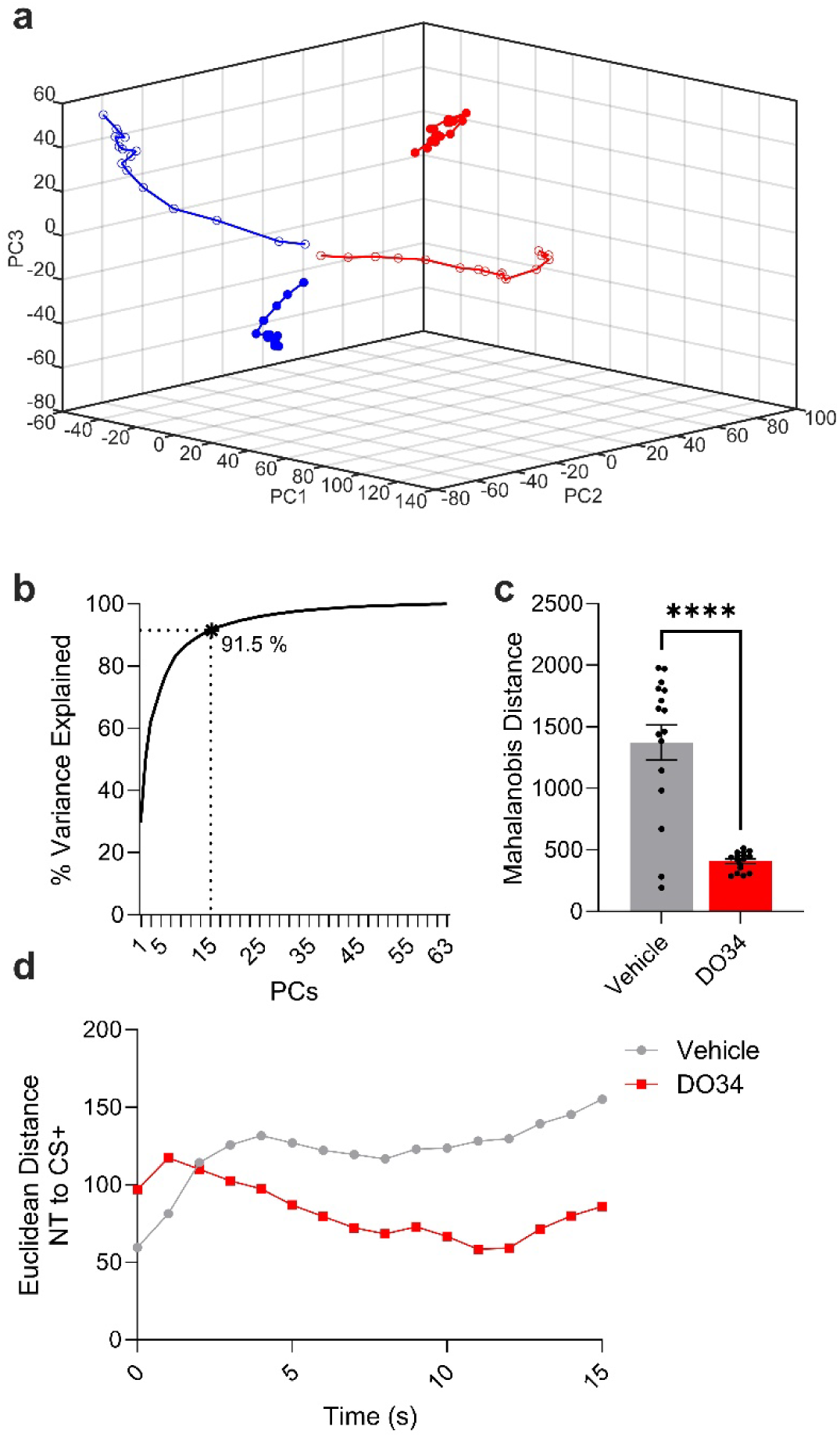
Reduced 2-AG is associated with increasing similarity of population metrics. (a) Trajectory of neuronal population activity in 3D principal component (PC) space during presentation of the CS+ (hollow dots) and NT (filled dots) for both DO34 (red) and vehicle-treated (blue) mice. (b) 15 PCs explain 91.5 % of the variance in the neuronal data set. (c) Mahalanobis distance between NT and CS+ trajectories in PC space for vehicle and DO34 exposed mice. (d) Instantaneous Euclidean distance between NT and CS+ trajectories across tone duration. Student’s t-test. ****p<0.0001.

### Rodent Data Analysis

Upon completion of experiments, freezing data was extracted from FreezeFrame. The time spent freezing to tones or context was expressed as a percentage of the interval. Freezing tones is shown in blocks of 2 tones. Statistical analysis and graphing were performed using GraphPad Prism 9 and the analysis performed is described in figure legends as appropriate.

Calcium imaging data was analyzed as previously described (Marcus et al., 2020; Patel et al., 2022). Briefly, data was preprocessed, bandpass filtered, and motion corrected using Inscopix Data Processing software V1.3. Individual neuron calcium traces were extracted using constrained non-negative matrix factorization for endoscopy algorithm (CNMFe)^35^ false positives were excluded from further analysis. The individual traces were aligned to tone onset and binned into 1s intervals. The binned traces were then averaged into blocks of 2 and normalized using Z-transform using the 15s pretone period as a normalization baseline. Traces exceeding a Z-score value of 3 or -3 (p = 0.01, two-tailed) for any 2 consecutive bins following tone onset during the tone period were considered (+) Responsive or (-) Responsive, respectively.

Agglomerative hierarchical clustering was conducted using the available MATLAB function clustergram. The clustergram function was applied to concatenated Z-scored traces for the NT and CS+. Linkage was then calculated using Ward’s method with Euclidean distance for metric. 3% of the maximum linkage was used as a threshold to form distinct clusters within the resulting dendrogram.

Principal component analysis (PCA) was conducted using available MATLAB functions. Z-scored traces for NT and CS+ periods were concatenated together for vehicle and DO34 groups. Concatenated traces for the DO34 group were randomly selected to create a matrix that had the same number of traces as the one for the vehicle group. The resulting vehicle and DO34 matrices were then concatenated and PCA was applied across the dimension of the activity traces. For visualization, the first 3 PCA dimensions for each group and stimulus type were plotted. To assess the distance between NT and CS+ in PC space, the 15d Mahalanobis distance between the NT and CS+ traces was calculated for vehicle and DO34 groups. To calculate the distance between NT and CS+ across the tone duration, we calculated the instantaneous Euclidean distance between the NT and CS+ for each 1s bin for the first 15 components in PC space.

### Data Availability

Data and custom codes used for analysis are available from the authors upon request.

## RESULTS

### Peripheral 2-AG levels are inversely correlated with conditioned fear generalization in humans

While preclinical studies clearly implicate 2-AG signaling in the regulation of stress-related behavioral phenotypes, clinical studies linking peripheral 2-AG levels to stress-related psychiatric diagnoses are mixed ^21,22,36–44^. To specifically examine the relationship between plasma 2-AG levels and fear learning, expression, and generalization we combined an experimental model of conditioning in human subjects together with measures of baseline plasma 2-AG (N=80) and resting state brain connectivity (N=88). Demographics of study subjects are shown in **Table 1**. **Figure 1a** depicts the experimental timeline and approach. On Day 1, subjects participated in screening session as described in the materials and methods. On Day 2, baseline 2-AG levels were obtained and subsequently, subjects were exposed to the fear conditioning protocol. On Day 3, a resting state imaging session was conducted. Using endocannabinoid levels as a co-variate in the fear conditioning analysis, we found a relationship between 2-AG and the ability to distinguish between the CS+, a stimulus predictive of an aversive outcome, and the CS-, a stimulus associated with the absence of an aversive outcome. Specifically, following acquisition there was a significant interaction between the startle response to CS-as compared to CS+ during fear recall (F1,78 = 4.10, p = 0.046, partial η2 = 0.05). Post-hoc linear regression with CS-vs. CS+ generalization [calculated as (CS-) – (CS+)] as dependent variable, and 2-AG levels as a predictor, confirmed that higher 2-AG predicted less generalization to CS-presentations (p = 0.025, 95% CI 0.007, 0.132, 1000 bootstrap samples; **Figure 1b**). Similarly, self-reported difficulties in emotion regulation (DERS) were associated with 2-AG levels such that lower peripheral levels of 2-AG predicted greater self-reported impairment in emotion regulation (r = -0.25, p = 0.021, N = 82; **Figure 1c**; F1,81 = 5.51, p = 0.013, 95% CI -27.0, -3.44, 1000 bootstraps).

### 2-AG depletion is associated with increased fear generalization in mice

Our human data indicate an inverse association between peripheral 2-AG levels and fear generalization, suggesting that lowering 2-AG levels could promote fear generalization. To test this hypothesis, we used a variety of fear conditioning paradigms in mice together with systemic injections of the DAGL inhibitor DO34 to pharmacologically inhibit the production of 2-AG prior to fear recall. Reducing 2-AG levels on Day 2 of a differential fear conditioning paradigm, prior to fear recall, did not increase freezing to CS- or CS+ presentation or generalization (**Figure 1d-f**). We hypothesized that our differential fear conditioning protocol in mice overtrained and minimized the ambiguity of the CS- or became associated as a safety cue which in turn minimized the effects of reducing 2-AG on fear generalization in mice. To test this hypothesis, we replicated our previous experiment, using a partial differential conditioning protocol where mice were presented with 5 CS-tones rather than 9 (**Figure 1g**). DO34 injections prior to fear recall increased freezing to CS-presentations (Treatment x tone interaction: F (3, 72) = 6.665, P=0.0005) and generalization (Treatment: F (1, 24) = 8.585, P=0.0073) without increasing freezing to CS+ presentation (**Figure 1g-i**). To further test the relationship between 2-AG levels and fear generalization, we used classical fear conditioning (**Figure 1j-l**), and on recall mice were presented with novel tones (NT) in addition to CS+ presentations. DO34 increases freezing to NTs (Treatment x tone interaction: F (3, 81) = 3.604, P=0.0169 and Treatment: F (1, 27) = 23.02, P<0.0001) and generalization (Treatment: F (1, 27) = 5.684, P=0.0244) without increasing freezing to CS+ presentation. We next expanded our studies to test the hypothesis that 2-AG is required for suppressing contextual fear generalization. Mice were conditioned in context A (CtxA) for 2 days and on Day 3, mice were injected with DO34 and exposed to CtxA or a novel context (NCtx; **Figure 1m-o**). Reducing 2-AG levels via DAGL inhibition increased freezing to NCTx without affecting freezing to CtxA (F (1, 33) = 11.03, P=0.0022) and increases generalization to NCtx (t(17)=2.320, p=0.0330). Our results suggest that 2-AG is required to for optimal control over generalization and that 2-AG depletion promotes fear generalization across multiple sensory modalities.

### Prefrontal cortical neuronal activity correlates of fear generalization during 2-AG deficient states

In rodents, PL has been implicated in fear expression and reduced generalization ^11,12,15^. However, how PL neurons discriminate between fear-predictive and non-predictive stimuli and how this process is affected by low 2-AG levels is not known. To address these questions, we conditioned mice to tones and the following day tested for CS+ recall and generalization to NTs after injections of vehicle or DO34 during *in vivo* single neuron calcium imaging sessions (**Figure 2a-c**). We recorded the activity of 802 PL neurons from 7 mice, and 885 neurons from 8 mice, treated with vehicle and DO34, respectively. DO34 increased net neuronal tone responses relative to pre-tone baseline in response to NT and CS+ (Main treatment effect: F (1, 77510) = 22.54, P<0.0001 and F (1, 77510) = 76.23, P<0.0001, respectively; **Figure 2d**). The increased net tone responses could be mediated by an increase in the magnitude of individual neuronal tone responses. To test this hypothesis, we identified neurons with tone responses exceeding +3 Z-scores ((+) Responsive, p = 0.01, two-tails) and those under -3 Z-scores ((-) Responsive) in any 2 consecutive 1s bins during the tone period and examined their respective average responses. As shown in **Figure 2e**, DO34 increased the magnitude of CS+ tone responses from (+) Responsive neurons (Treatment x trial interaction: F (45, 18270) = 1.542, P=0.0114) without changing the magnitude of the responses from (-) Responsive neurons. Thus, the increased (+) Responsive magnitude appears to explain the increased net response to CS+ tones. Interestingly, DO34 lead to a quicker return to baseline from (-) Responsive neurons (Treatment x trial interaction: F (45, 7560) = 2.050, P<0.0001 and main treatment effect: F (1, 168) = 4.945, P=0.0275) without impacting the magnitude of (+) responses.

A larger proportion of tone responsive neurons could be an important contributor to the higher net neuronal activity to tones seen with DO34. To gain some insight into the diversity of neuronal responses to distinct tone types, we conducted agglomerative hierarchical clustering of neuronal tone responses. We observed clusters of neurons that do not respond to tones, neurons that exclusively respond to one tone type, and neurons with diverse types of dual responses to tones with the DO34 group grossly having a larger proportion of dual responses (**Figure 2f-g**). We then formally quantified the proportion of different neuronal tone response types. We found that DO34 increased the proportion of overall neuronal tone responses (Fisher’s Exact Test, p<0.00001; **Figure 2h-i**). Most importantly, DO34 increased the proportion of dual tone responsive neurons (Fisher’s Exact Test, p<0.0052) and decreased the proportion of CS+ (+) tone responses (Fisher’s Exact Test, p<0.0093) without impacting the proportion of NT responsive neurons. The profile of dual responsive neurons was heterogenous as shown in **Figure S1**. The observed increase in tone responsivity is not simply explained by increased neuronal excitability but by increased likelihood of responding specifically to presentation of tones, as we actually observed a small reduction in spontaneous calcium events after DO34 treatment (H(3)= 47.53, p <0.0001; **Figure S2**). In addition, DO34-treated mice showed a greater proportion of neurons that responded to consecutive tones of any type (Fisher’s Exact Test: p< 0.00001 and p<0.00001 for NT and CS+ respectively; **Figure S3**), suggesting that 2-AG depletion increases ensemble stability in response to sensory stimuli presentation. Our results suggest that generalization is associated with an increase in dual-responsive PL neurons that are typically specific to CS+, while the proportion of NT active neurons remains stable.

Upon analysis of neurons longitudinally registered between conditioning and recall, we found that neuronal tone or shock responsiveness during conditioning did not predict tone responsiveness on recall as hierarchical agglomerative clustering only revealed a small cluster of tone/shock responsive neurons that also responded to tones during the recall (Figure **S4**). Moreover, only ∼4% of neurons that responded to CS+ during the late conditioning phase responded to NTs or CS+ during recall. These data suggest a high degree of ensemble drift across conditioning and recall with respect to tone responsive neurons in PL. This finding contrasts with evidence that amygdala ensembles are stable across days following fear conditioning ^45^.

Our data above indicating an increase in dual-responsive neurons after 2-AG depletion suggest that DO34 could increase generalization by increasing the similarity of the population activity associated with NT and CS+ presentations. To address this hypothesis, we used a population vector approach and PCA for dimensionality reduction of neuronal activity during tone presentations. **Figure 3a** depicts the trajectory of population activity in 3D PC space during presentation of the CS+ and NT for both DO34 and vehicle-treated mice. We used 15 principal components (PC), which accounted for 91.5 % of the variance in our data set, for our analysis (**Figure 3b**). We then formally quantified the distance between NT and CS+ in PC space. As shown in **Figure 3c**, DO34 decreased the Mahalanobis distance between NT and CS+ activity in PC space relative to vehicle treatment (t(30)= t=6.705, p <0.0001). Moreover, despite starting higher, the Euclidean distance between NT and CS+ activity trajectories decreased across tone presentations in DO34 treated mice, while the activity distance for the vehicle group continues to rise (**Figure 3d**). These results suggest that DO34 increases the similarity of population activity dynamics between NT and CS+, which could represent an important neural correlate of fear generalization.

### Prefrontal-Amygdala Connectivity is inversely correlates with fear generalization

Our results implicate the PFC in fear generalization/discrimination and Amygdala-PFC connectivity has been implicated in fear expression and reduced generalization ^11,15,46,47^. To explore if fear generalization and 2-AG levels are associated with amygdala-PFC functional connectivity, we obtained and analyzed brain resting state data. Beta coefficients for right/left amygdala were assessed for covariance with peripheral 2-AG levels and behavioral measures of fear conditioning. As shown in **Figure 4d**, CS generalization explains decreased connectivity between the right amygdala and dorsolateral prefrontal cortex (dlPFC). Specifically, we identified a positive cluster in dlPFC (area 46, MNI: -44, 37, 28, 5 voxels), with CS generalization as a covariate. Post-hoc linear regression on extracted ß-coefficients as the dependent variable and CS generalization as the predictor confirmed the voxel-based results (F1,87 = 16.3; p < 0.001; 95% CI [0.022, 0.051]; 1000 boostrap samples; **Figure 4e**). We did not observe any significant association between 2-AG levels and amygdala resting state connectivity or between dorsal anterior cingulate cortex and amygdala resting state connectivity.

**Fig. 4.**
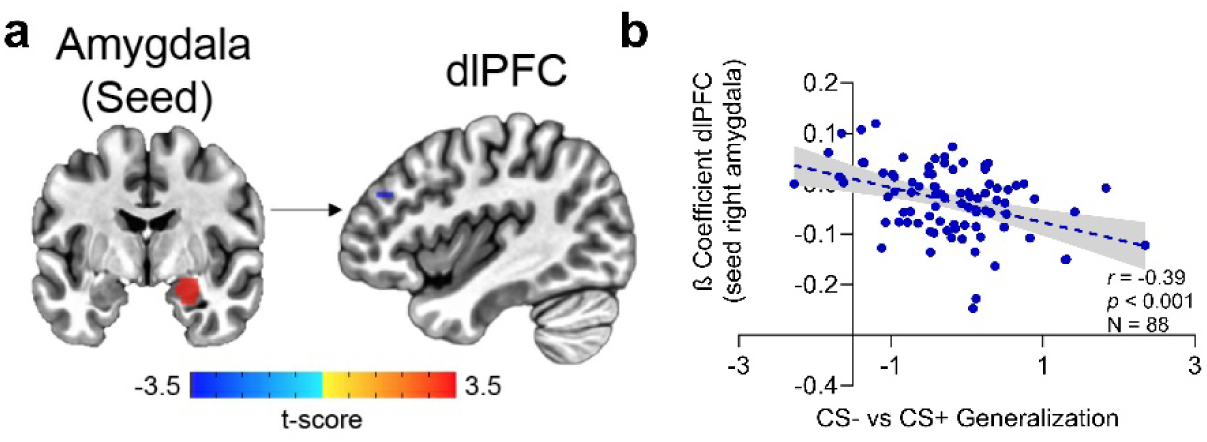
dlPFC-amygdala resting state connectivity is inversely correlated to fear generalization. (a) Depiction of increased resting-state between the right amygdala and dlPFC. (b) β coefficient for dlPFC as a function of generalization between CS- and CS+.

## DISCUSSION

Using distinct but complementary approaches we show, from mice to humans, that 2-AG regulates the balance between fear discrimination and generalization. Plasma levels of 2-AG were inversely correlated with generalization in humans and pharmacological 2-AG depletion increased fear generalization in mice using diverse behavioral paradigms. Moreover, this increased generalization was associated with impaired sensory discrimination within PL neuronal ensembles responsive to sensory cues.

We show that plasma levels of 2-AG predict discriminatory performance without impacting CS+ responses. There are 3 potential explanations for this: 1) Spillover of tonic neurotransmitter release could have predictive value over subsequent task performance; 2) peripherally produced 2-AG permeates across the blood-brain barrier and into neuronal synapses which in turn has a modulatory effect over behavior; or 3) plasma 2-AG levels reflect conserved (central-peripheral) within-subject enzymatic balance between production and degradation of 2-AG that varies depending on individual environmental, developmental, genetic, and epigenetic factors. Of these, the first possibility is less likely as knockout mice for DAGLα have normal levels of plasma 2-AG despite having low brain 2-AG levels, thus, brain levels seem not to have an effect over plasma levels ^17^. The second possibility is also less likely, as synaptic 2-AG signaling is tightly regulated by local synthesis and degradation on the time scale of seconds, which is unlikely to be affected by relatively slow changes in circulating 2-AG. Importantly, subjects used in this study had a differential history of trauma and substance use. Indeed, this heterogeneity may be required for observing a relationship between 2-AG and fear generalization as this was not observed during analysis of subject subsets. This may provide further support for the third hypothesis noted above. Previous studies examining peripheral 2-AG levels and PTSD symptoms have been mixed ^21,22,38–41,43^, wherein some studies have shown an association between lower levels of 2-AG and PTSD diagnosis after traumatic stress exposure ^21,41^. In contrast, higher 2-AG levels immediately post trauma exposure was associated with subsequent PTSD diagnosis and positively correlated with symptom severity in racial minority subjects ^48^. Future studies aimed at more clearly defining the relationship between peripheral and central 2-AG levels and between 2-AG levels and aspects of trauma and PTSD will be critical to resolve these apparent discrepancies in the literature.

The dorsolateral PFC (dlPFC) has been implicated in emotional regulation, acquisition and expression of fear, and reduced generalization/improved discrimination ^49,50^. Our results further implicate the prefrontal cortex in discrimination between safety and danger, as prefrontal-amygdala connectivity was predictive of discrimination between safe and aversive stimuli. Intriguingly, we found an association between dlPFC-amygdala, but not dACC-amygdala, connectivity and generalization. In part, this finding could be related to the fact that our data was obtained during resting state and studies implicating dACC in fear expression and functional homology to the rodent PL characterize dACC activity in response to cue presentations ^51–53^. The dlPFC does not directly connect with the amygdala and it has been proposed that dlPFC exerts cognitive control over amygdala activity through the ventromedial prefrontal cortex ^54–56^. The dlPFC also targets the dACC ^57,58^ and in turn dACC targets the amygdala ^59,60^; we suggest that dlPFC-dACC-amygdala circuit could provide an important route to exert control over amygdala neutral stimuli responses and reduce fear generalization. Our findings implicating dlPFC-amygdala resting state connectivity and decreased generalization may suggest that resting state connectivity could be an intrinsic indicator of potential entrainment strength of dlPFC over amygdala to suppress generalized threat responses. These data also suggest that resting state functional connectivity between PFC and amygdala could be useful as a biomarker with predictive value for fear generalization or even development of PTSD, though confirmatory studies are warranted. One potential limitation of our study is that the imaging session was conducted on a separate day from the fear conditioning, thus it is unclear if resting connectivity would be different from that obtained prior to or during fear conditioning.

In rodents, the ventromedial prefrontal cortex has been implicated in reducing fear generalization^11^. Specifically, the infralimbic prefrontal cortex regulates safety learning and reduction of conditioned fear and generalization ^8,61–65^. The prelimbic cortex mediates conditioned fear expression, safety learning, and reduction of fear generalization ^8,12–15,66–68^. Our results show that PL contains information needed for reducing generalization as loss of specificity in PL CS+ neuronal ensembles promoted fear generalization. This idea is supported by another study indicating the overlap (and associated behavioral generalization) between PL neuronal subpopulation activated by CS+ and NT increases with increasing frequency similarity between the two stimuli ^69^. Thus, neuronal ensembles within PL simultaneously representing both auditory sensory stimuli, and fear generalization may be melding together of both populations and concurrent increased similarity between NT and CS+ population dynamics.

Previous studies in rodents have shown sex differences in fear generalization and expression of other conditioned responses ^70–72^. Our study did not find any sex differences in generalization or 2-AG measures in human subjects. In part this difference could be attributed to conditioning protocol used, interspecies differences, or a modest sample size that may preclude adequate group sizes to formally test for such differences. Despite the lack of sex differences in humans, one limitation of our study is that we only examined the behavior and neuronal activity of male mice.

Our study provides mechanistic insight into how fear generalization emerges in the prefrontal cortex and how this is controlled by 2-AG signaling. These data provide cross-species evidence that 2-AG plays a critical role in opposing fear generalization and support the notion that 2-AG deficient states may contribute to the development of trauma-related disorders including PTSD. Intriguingly, our results suggest that plasma 2-AG levels and dlPFC-amygdala connectivity could be independent biomarkers predicting the degree of fear generalization. Future studies may focus on assessing if these features, at least on the extremes of very low or high states, are predictive of resilience or susceptibility to trauma.

**Disclosures:** Dr. Patel is a scientific consultant for Psy Therapeutics, Janssen Pharmaceuticals, and Jazz Pharmaceuticals for work unrelated to the topic of this manuscript. Dr. Heilig has received consulting fees, research support, or other compensation from Indivior, Camurus, BrainsWay, Aelis Farma, and Janssen Pharmaceuticals. Dr. Mayo is a scientific consultant for Synendos Therapeutics. The other authors report no financial relationships with commercial interests.

## Acknowledgements

Research funding from NIMH grants K08MH126166 to Dr. Rosas-Vidal and R01MH107435 to Dr. Sachin Patel. Research funding from NARSAD Young Investigator Grants from the Brain & Behavior Research Foundation #29255 to Dr. Rosas-Vidal and #27094 to Dr. Mayo. Dr. Heilig is supported by the Swedish Research Council grants 2013-07434, 2019-01138 and is a Wallenberg Foundation Clinical Scholar. We thank Edmund Havener for technical assistance.

## References

1. Lissek S, Powers AS, McClure EB, et al. Classical fear conditioning in the anxiety disorders: a meta-analysis. Behav Res Ther. Nov 2005;43(11):1391–424. doi:10.1016/j.brat.2004.10.007

2. Lissek S, Biggs AL, Rabin SJ, et al. Generalization of conditioned fear-potentiated startle in humans: experimental validation and clinical relevance. Behav Res Ther. May 2008;46(5):678–87. doi:10.1016/j.brat.2008.02.005

3. Lissek S, Bradford DE, Alvarez RP, et al. Neural substrates of classically conditioned fear-generalization in humans: a parametric fMRI study. Soc Cogn Affect Neurosci. Aug 2014;9(8):1134–42. doi:10.1093/scan/nst096

4. Lissek S, Rabin S, Heller RE, et al. Overgeneralization of conditioned fear as a pathogenic marker of panic disorder. Am J Psychiatry. Jan 2010;167(1):47–55. doi:10.1176/appi.ajp.2009.09030410

5. Morey RA, Dunsmoor JE, Haswell CC, et al. Fear learning circuitry is biased toward generalization of fear associations in posttraumatic stress disorder. Transl Psychiatry. Dec 15 2015;5(12):e700. doi:10.1038/tp.2015.196

6. Ghosh S, Chattarji S. Neuronal encoding of the switch from specific to generalized fear. Nat Neurosci. Jan 2015;18(1):112–20. doi:10.1038/nn.3888

7. Onat S, Buchel C. The neuronal basis of fear generalization in humans. Nat Neurosci. Dec 2015;18(12):1811–8. doi:10.1038/nn.4166

8. Corches A, Hiroto A, Bailey TW, et al. Differential fear conditioning generates prefrontal neural ensembles of safety signals. Behav Brain Res. Mar 15 2019;360:169–184. doi:10.1016/j.bbr.2018.11.042

9. Antoniadis EA, McDonald RJ. Fornix, medial prefrontal cortex, nucleus accumbens, and mediodorsal thalamic nucleus: roles in a fear-based context discrimination task. Neurobiol Learn Mem. Jan 2006;85(1):71–85. doi:10.1016/j.nlm.2005.08.011

10. Xu W, Sudhof TC. A neural circuit for memory specificity and generalization. Science. Mar 15 2013;339(6125):1290–5. doi:10.1126/science.1229534

11. Likhtik E, Stujenske JM, Topiwala MA, Harris AZ, Gordon JA. Prefrontal entrainment of amygdala activity signals safety in learned fear and innate anxiety. Nat Neurosci. Jan 2014;17(1):106–13. doi:10.1038/nn.3582

12. Burgos-Robles A, Vidal-Gonzalez I, Quirk GJ. Sustained conditioned responses in prelimbic prefrontal neurons are correlated with fear expression and extinction failure. J Neurosci. Jul 1 2009;29(26):8474–82. doi:10.1523/JNEUROSCI.0378-09.2009

13. Sierra-Mercado D, Padilla-Coreano N, Quirk GJ. Dissociable roles of prelimbic and infralimbic cortices, ventral hippocampus, and basolateral amygdala in the expression and extinction of conditioned fear. Neuropsychopharmacology. Jan 2011;36(2):529–38. doi:10.1038/npp.2010.184

14. Rozeske RR, Jercog D, Karalis N, et al. Prefrontal-Periaqueductal Gray-Projecting Neurons Mediate Context Fear Discrimination. Neuron. Feb 21 2018;97(4):898–910 e6. doi:10.1016/j.neuron.2017.12.044

15. Stujenske JM, O’Neill PK, Fernandes-Henriques C, et al. Prelimbic cortex drives discrimination of non-aversion via amygdala somatostatin interneurons. Neuron. Jul 20 2022;110(14):2258–2267 e11. doi:10.1016/j.neuron.2022.03.020

16. Scheyer A, Yasmin F, Naskar S, Patel S. Endocannabinoids at the synapse and beyond: implications for neuropsychiatric disease pathophysiology and treatment. Neuropsychopharmacology. Jan 2023;48(1):37–53. doi:10.1038/s41386-022-01438-7

17. Shonesy BC, Bluett RJ, Ramikie TS, et al. Genetic disruption of 2-arachidonoylglycerol synthesis reveals a key role for endocannabinoid signaling in anxiety modulation. Cell Rep. Dec 11 2014;9(5):1644–1653. doi:10.1016/j.celrep.2014.11.001

18. Lutz B, Marsicano G, Maldonado R, Hillard CJ. The endocannabinoid system in guarding against fear, anxiety and stress. Nat Rev Neurosci. Dec 2015;16(12):705–18. doi:10.1038/nrn4036

19. Bluett RJ, Baldi R, Haymer A, et al. Endocannabinoid signalling modulates susceptibility to traumatic stress exposure. Nat Commun. Mar 28 2017;8:14782. doi:10.1038/ncomms14782

20. Cavener VS, Gaulden A, Pennipede D, et al. Inhibition of Diacylglycerol Lipase Impairs Fear Extinction in Mice. Front Neurosci. 2018;12:479. doi:10.3389/fnins.2018.00479

21. Hill MN, Bierer LM, Makotkine I, et al. Reductions in circulating endocannabinoid levels in individuals with post-traumatic stress disorder following exposure to the World Trade Center attacks. Psychoneuroendocrinology. Dec 2013;38(12):2952–61. doi:10.1016/j.psyneuen.2013.08.004

22. Leen NA, de Weijer AD, van Rooij SJH, Kennis M, Baas JMP, Geuze E. The Role of the Endocannabinoids 2-AG and Anandamide in Clinical Symptoms and Treatment Outcome in Veterans with PTSD. Chronic Stress (Thousand Oaks). Jan-Dec 2022;6:24705470221107290. doi:10.1177/24705470221107290

23. Bjureberg J, Ljotsson B, Tull MT, et al. Development and Validation of a Brief Version of the Difficulties in Emotion Regulation Scale: The DERS-16. J Psychopathol Behav Assess. Jun 2016;38(2):284–296. doi:10.1007/s10862-015-9514-x

24. Perini I, Mayo LM, Capusan AJ, et al. Resilience to substance use disorder following childhood maltreatment: association with peripheral biomarkers of endocannabinoid function and neural indices of emotion regulation. Mol Psychiatry. Apr 12 2023;doi:10.1038/s41380-023-02033-y

25. Capusan AJ, Gustafsson PA, Kuja-Halkola R, Igelstrom K, Mayo LM, Heilig M. Re-examining the link between childhood maltreatment and substance use disorder: a prospective, genetically informative study. Mol Psychiatry. Jul 2021;26(7):3201–3209. doi:10.1038/s41380-021-01071-8

26. Mayo LM, Asratian A, Linde J, et al. Protective effects of elevated anandamide on stress and fear-related behaviors: translational evidence from humans and mice. Mol Psychiatry. May 2020;25(5):993–1005. doi:10.1038/s41380-018-0215-1

27. Mayo LM, Asratian A, Linde J, et al. Elevated Anandamide, Enhanced Recall of Fear Extinction, and Attenuated Stress Responses Following Inhibition of Fatty Acid Amide Hydrolase: A Randomized, Controlled Experimental Medicine Trial. Biol Psychiatry. Mar 15 2020;87(6):538–547. doi:10.1016/j.biopsych.2019.07.034

28. Milad MR, Orr SP, Pitman RK, Rauch SL. Context modulation of memory for fear extinction in humans. Psychophysiology. Jul 2005;42(4):456–64. doi:10.1111/j.1469-8986.2005.00302.x

29. Neumann DL, Waters AM. The use of an unpleasant sound as an unconditional stimulus in a human aversive Pavlovian conditioning procedure. Biol Psychol. Aug 2006;73(2):175–85. doi:10.1016/j.biopsycho.2006.03.004

30. Walker DL, Davis M. Quantifying fear potentiated startle using absolute versus proportional increase scoring methods: implications for the neurocircuitry of fear and anxiety. Psychopharmacology (Berl). Nov 2002;164(3):318–28. doi:10.1007/s00213-002-1213-0

31. Blumenthal TD, Cuthbert BN, Filion DL, Hackley S, Lipp OV, van Boxtel A. Committee report: Guidelines for human startle eyeblink electromyographic studies. Psychophysiology. Jan 2005;42(1):1–15. doi:10.1111/j.1469-8986.2005.00271.x

32. Stensson N, Ghafouri B, Gerdle B, Ghafouri N. Alterations of anti-inflammatory lipids in plasma from women with chronic widespread pain - a case control study. Lipids Health Dis. Jun 12 2017;16(1):112. doi:10.1186/s12944-017-0505-7

33. Cox RW. AFNI: software for analysis and visualization of functional magnetic resonance neuroimages. Comput Biomed Res. Jun 1996;29(3):162–73. doi:10.1006/cbmr.1996.0014

34. Cox RW, Chen G, Glen DR, Reynolds RC, Taylor PA. FMRI Clustering in AFNI: False-Positive Rates Redux. Brain Connect. Apr 2017;7(3):152–171. doi:10.1089/brain.2016.0475

35. Zhou P, Resendez SL, Rodriguez-Romaguera J, et al. Efficient and accurate extraction of in vivo calcium signals from microendoscopic video data. Elife. Feb 22 2018;7doi:10.7554/eLife.28728

36. Hill MN, Miller GE, Carrier EJ, Gorzalka BB, Hillard CJ. Circulating endocannabinoids and N-acyl ethanolamines are differentially regulated in major depression and following exposure to social stress. Psychoneuroendocrinology. Sep 2009;34(8):1257–62. doi:10.1016/j.psyneuen.2009.03.013

37. Hill MN, Miller GE, Ho WS, Gorzalka BB, Hillard CJ. Serum endocannabinoid content is altered in females with depressive disorders: a preliminary report. Pharmacopsychiatry. Mar 2008;41(2):48–53. doi:10.1055/s-2007-993211

38. Hauer D, Schelling G, Gola H, et al. Plasma concentrations of endocannabinoids and related primary fatty acid amides in patients with post-traumatic stress disorder. PLoS One. 2013;8(5):e62741. doi:10.1371/journal.pone.0062741

39. Neumeister A, Normandin MD, Pietrzak RH, et al. Elevated brain cannabinoid CB1 receptor availability in post-traumatic stress disorder: a positron emission tomography study. Mol Psychiatry. Sep 2013;18(9):1034–40. doi:10.1038/mp.2013.61

40. Schaefer C, Enning F, Mueller JK, et al. Fatty acid ethanolamide levels are altered in borderline personality and complex posttraumatic stress disorders. Eur Arch Psychiatry Clin Neurosci. Aug 2014;264(5):459–63. doi:10.1007/s00406-013-0470-8

41. Crombie KM, Leitzelar BN, Brellenthin AG, Hillard CJ, Koltyn KF. Loss of exercise- and stress-induced increases in circulating 2-arachidonoylglycerol concentrations in adults with chronic PTSD. Biol Psychol. Jul 2019;145:1–7. doi:10.1016/j.biopsycho.2019.04.002

42. Fitzgerald JM, Chesney SA, Lee TS, et al. Circulating endocannabinoids and prospective risk for depression in trauma-injury survivors. Neurobiol Stress. May 2021;14:100304. doi:10.1016/j.ynstr.2021.100304

43. Botsford C, Brellenthin AG, Cisler JM, Hillard CJ, Koltyn KF, Crombie KM. Circulating endocannabinoids and psychological outcomes in women with PTSD. J Anxiety Disord. Jan 2023;93:102656. doi:10.1016/j.janxdis.2022.102656

44. Walther A, Kirschbaum C, Wehrli S, et al. Depressive symptoms are negatively associated with hair N-arachidonoylethanolamine (anandamide) levels: A cross-lagged panel analysis of four annual assessment waves examining hair endocannabinoids and cortisol. Prog Neuropsychopharmacol Biol Psychiatry. Mar 8 2023;121:110658. doi:10.1016/j.pnpbp.2022.110658

45. Grewe BF, Grundemann J, Kitch LJ, et al. Neural ensemble dynamics underlying a long-term associative memory. Nature. Mar 30 2017;543(7647):670–675. doi:10.1038/nature21682

46. Fani N, King TZ, Brewster R, et al. Fear-potentiated startle during extinction is associated with white matter microstructure and functional connectivity. Cortex. Mar 2015;64:249–59. doi:10.1016/j.cortex.2014.11.006

47. Greco JA, Liberzon I. Neuroimaging of Fear-Associated Learning. Neuropsychopharmacology. Jan 2016;41(1):320–34. doi:10.1038/npp.2015.255

48. deRoon-Cassini TA, Bergner CL, Chesney SA, et al. Circulating endocannabinoids and genetic polymorphisms as predictors of posttraumatic stress disorder symptom severity: heterogeneity in a community-based cohort. Transl Psychiatry. Feb 1 2022;12(1):48. doi:10.1038/s41398-022-01808-1

49. LaBar KS, Cabeza R. Cognitive neuroscience of emotional memory. Nat Rev Neurosci. Jan 2006;7(1):54–64. doi:10.1038/nrn1825

50. Lam CLM, Wong CHY, Junghofer M, Roesmann K. Implicit threat learning involves the dorsolateral prefrontal cortex and the cerebellum. Int J Clin Health Psychol. Apr-Jun 2023;23(2):100357. doi:10.1016/j.ijchp.2022.100357

51. Milad MR, Quirk GJ, Pitman RK, Orr SP, Fischl B, Rauch SL. A role for the human dorsal anterior cingulate cortex in fear expression. Biol Psychiatry. Nov 15 2007;62(10):1191–4. doi:10.1016/j.biopsych.2007.04.032

52. Fullana MA, Harrison BJ, Soriano-Mas C, et al. Neural signatures of human fear conditioning: an updated and extended meta-analysis of fMRI studies. Mol Psychiatry. Apr 2016;21(4):500–8. doi:10.1038/mp.2015.88

53. Rougemont-Bucking A, Linnman C, Zeffiro TA, et al. Altered processing of contextual information during fear extinction in PTSD: an fMRI study. CNS Neurosci Ther. Aug 2011;17(4):227–36. doi:10.1111/j.1755-5949.2010.00152.x

54. Kroes MCW, Dunsmoor JE, Hakimi M, et al. Patients with dorsolateral prefrontal cortex lesions are capable of discriminatory threat learning but appear impaired in cognitive regulation of subjective fear. Soc Cogn Affect Neurosci. Aug 7 2019;14(6):601–612. doi:10.1093/scan/nsz039

55. Alexandra Kredlow M, Fenster RJ, Laurent ES, Ressler KJ, Phelps EA. Prefrontal cortex, amygdala, and threat processing: implications for PTSD. Neuropsychopharmacology. Jan 2022;47(1):247–259. doi:10.1038/s41386-021-01155-7

56. Ochsner KN, Gross JJ. The cognitive control of emotion. Trends Cogn Sci. May 2005;9(5):242–9. doi:10.1016/j.tics.2005.03.010

57. Barbas H, Pandya DN. Architecture and intrinsic connections of the prefrontal cortex in the rhesus monkey. J Comp Neurol. Aug 15 1989;286(3):353–75. doi:10.1002/cne.902860306

58. Selemon LD, Goldman-Rakic PS. Common cortical and subcortical targets of the dorsolateral prefrontal and posterior parietal cortices in the rhesus monkey: evidence for a distributed neural network subserving spatially guided behavior. J Neurosci. Nov 1988;8(11):4049–68. doi:10.1523/JNEUROSCI.08-11-04049.1988

59. Stevens FL, Hurley RA, Taber KH. Anterior cingulate cortex: unique role in cognition and emotion. J Neuropsychiatry Clin Neurosci. Spring 2011;23(2):121–5. doi:10.1176/jnp.23.2.jnp121

60. Zikopoulos B, Hoistad M, John Y, Barbas H. Posterior Orbitofrontal and Anterior Cingulate Pathways to the Amygdala Target Inhibitory and Excitatory Systems with Opposite Functions. J Neurosci. May 17 2017;37(20):5051–5064. doi:10.1523/JNEUROSCI.3940-16.2017

61. Bravo-Rivera C, Roman-Ortiz C, Brignoni-Perez E, Sotres-Bayon F, Quirk GJ. Neural structures mediating expression and extinction of platform-mediated avoidance. J Neurosci. Jul 16 2014;34(29):9736–42. doi:10.1523/JNEUROSCI.0191-14.2014

62. Sangha S, Robinson PD, Greba Q, Davies DA, Howland JG. Alterations in reward, fear and safety cue discrimination after inactivation of the rat prelimbic and infralimbic cortices. Neuropsychopharmacology. Sep 2014;39(10):2405–13. doi:10.1038/npp.2014.89

63. Do-Monte FH, Manzano-Nieves G, Quinones-Laracuente K, Ramos-Medina L, Quirk GJ. Revisiting the role of infralimbic cortex in fear extinction with optogenetics. J Neurosci. Feb 25 2015;35(8):3607–15. doi:10.1523/JNEUROSCI.3137-14.2015

64. Felix-Ortiz AC, Terrell JM, Gonzalez C, et al. The infralimbic and prelimbic cortical areas bidirectionally regulate safety learning during normal and stress conditions. bioRxiv. May 5 2023;doi:10.1101/2023.05.05.539516

65. Ng K, Pollock M, Escobedo A, et al. Suppressing fear in the presence of a safety cue requires infralimbic cortical signaling to central amygdala. Neuropsychopharmacology. May 15 2023;doi:10.1038/s41386-023-01598-0

66. Rosas-Vidal LE, Lozada-Miranda V, Cantres-Rosario Y, Vega-Medina A, Melendez L, Quirk GJ. Alteration of BDNF in the medial prefrontal cortex and the ventral hippocampus impairs extinction of avoidance. Neuropsychopharmacology. Dec 2018;43(13):2636–2644. doi:10.1038/s41386-018-0176-8

67. Sangha S, Diehl MM, Bergstrom HC, Drew MR. Know safety, no fear. Neurosci Biobehav Rev. Jan 2020;108:218-230. doi:10.1016/j.neubiorev.2019.11.006

68. Jercog D, Winke N, Sung K, et al. Dynamical prefrontal population coding during defensive behaviours. Nature. Jul 2021;595(7869):690–694. doi:10.1038/s41586-021-03726-6

69. Grosso A, Santoni G, Manassero E, Renna A, Sacchetti B. A neuronal basis for fear discrimination in the lateral amygdala. Nat Commun. Mar 23 2018;9(1):1214. doi:10.1038/s41467-018-03682-2

70. Gruene TM, Flick K, Stefano A, Shea SD, Shansky RM. Sexually divergent expression of active and passive conditioned fear responses in rats. Elife. Nov 14 2015;4 doi:10.7554/eLife.11352

71. Keiser AA, Turnbull LM, Darian MA, Feldman DE, Song I, Tronson NC. Sex Differences in Context Fear Generalization and Recruitment of Hippocampus and Amygdala during Retrieval. Neuropsychopharmacology. Jan 2017;42(2):397–407. doi:10.1038/npp.2016.174

72. Trask S, Reis DS, Ferrara NC, Helmstetter FJ. Decreased cued fear discrimination learning in female rats as a function of estrous phase. Learn Mem. Jun 2020;27(6):254–257. doi:10.1101/lm.051185.119

