## Supplementary Figures and Methods for "PREFRONTAL CORRELATES OF FEAR GENERALIZATION DURING ENDOCANNABINOID DEPLETION"

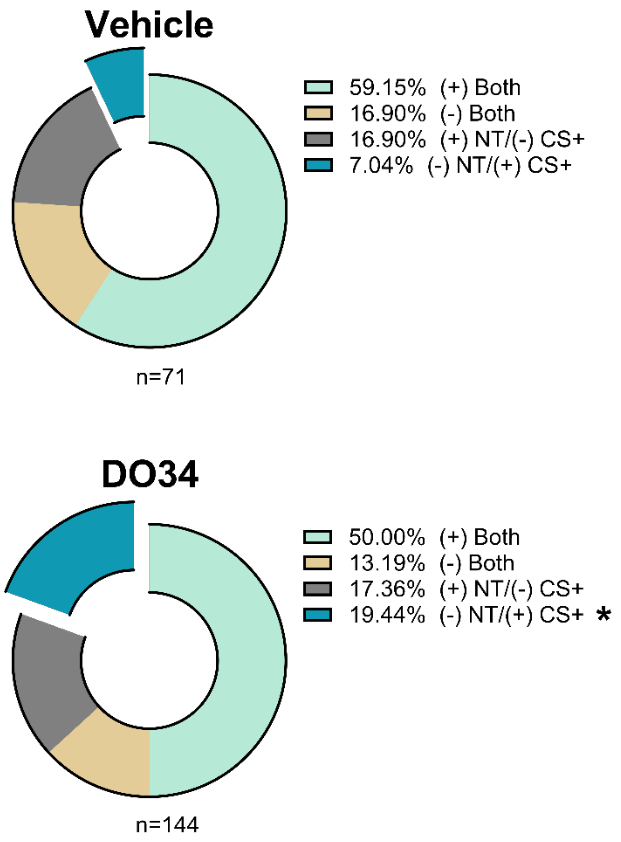


**S1. Diversity of Dual responses.** Proportion of neurons that respond significantly to both NT and CS+ by increasing their activity to both NT and CS+ ((+) Both), decreasing their activity to both NT and CS+ ((-)Both), increasing their activity to NT and decreasing their activity to CS+ ((+)NT/(-)CS+), and decreasing their activity to NT and increasing their activity to CS+ ((-)NT/(+)CS+) from vehicle (Top) and DO34 (bottom) exposed mice. Fisher's Exact Test: *$p < 0.05$.

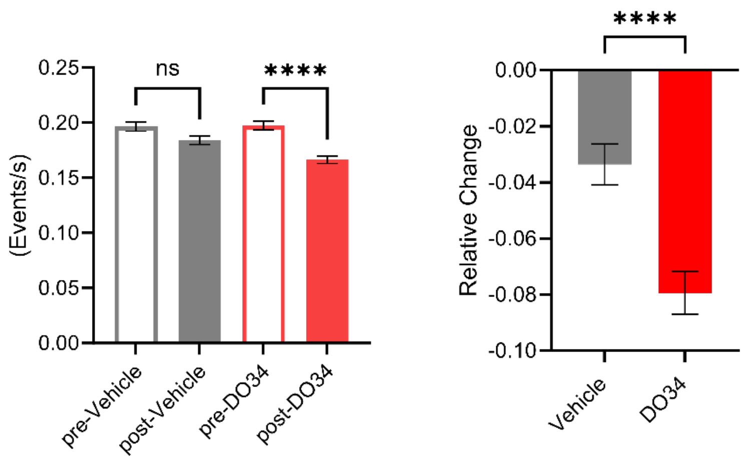


**S2. Spontaneous calcium event rate.** Left. Calcium transient event rate before and after vehicle or DO34 treatment. Right. Relative change in calcium event rate following treatment. Two-way Kruskal-Wallis and Mann-Whitney test as appropriate. ****p<0.0001.

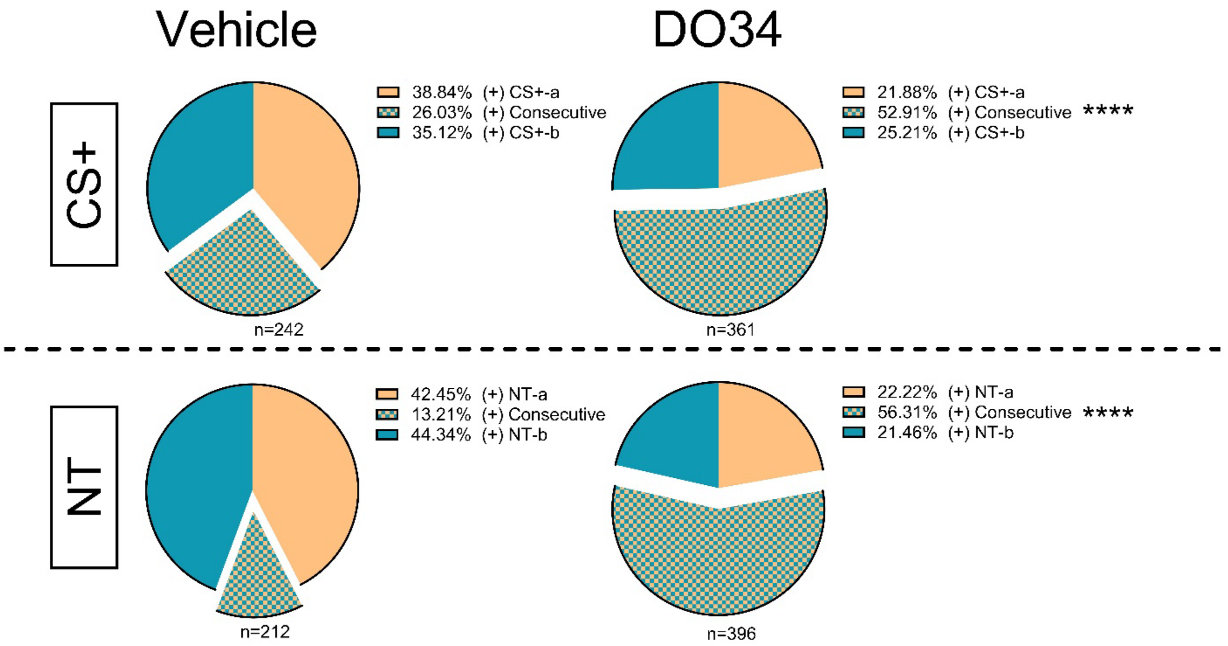


**S3. DO34 increases the proportion of neurons that respond consecutively to tones of the same type.** Top. Proportion of neurons that respond positively to the first CS+ presentation out of a consecutive pair ((+)CS+-a), to the 2 consecutive CS+ presentations ((+)Consecutive), or the last CS+ presentation of the pair ((+)CS+-b; from Vehicle (left) and DO34 (right) exposed mice. Bottom. Proportion of neurons that respond positively to the first NT presentation out of a consecutive pair ((+)NT-a), to the 2 consecutive NT presentations ((+)Consecutive), or the last NT presentation of the pair ((+)NT-b; from Vehicle (left) and DO34 (right) exposed mice. Fisher's exact test. ****p<0.0001..

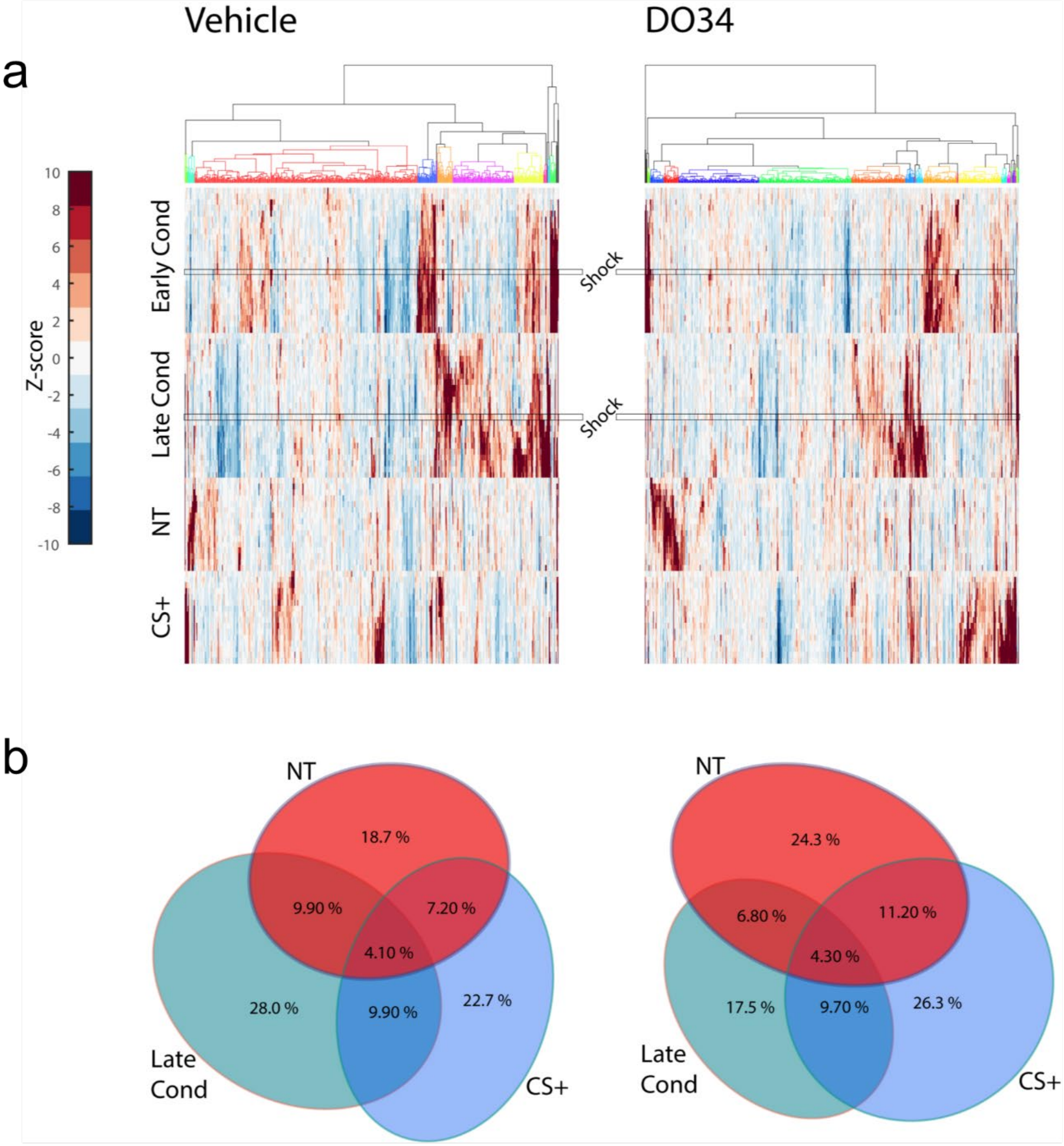


**S4. Neuronal tone responses or shock responses do not predict tone responsivity the following day.** (a) Hierarchical tree clustering of Z-scored neuronal activity to CS+, foot shock (demarcated by horizontal rectangle), and NT presentations during early and late conditioning phases and recall phases from vehicle (left) and DO34 (right) exposed mice. (b) Venn diagrams of overlap between the proportion of neurons that respond to late conditioning CS+ and NT and CS+ during recall phase the following day from vehicle (left) and DO34 (right) exposed mice.

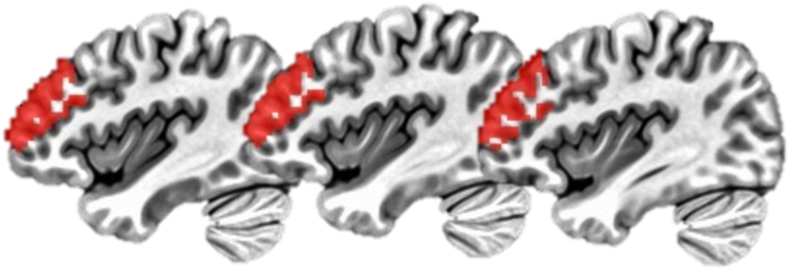

**Figure S5**. Mask used for the resting-state functional connectivity analysis, superimposed on a standard MNI template. The mask included the dlPFC region, defined by the Matthew Glasser atlas.

### Materials and methods

#### Human Participants

Participants were recruited between March 2017 and July 2020, at Linköping University. A total of 101 participants were included in the study, divided into four groups across the dimensions of histories of childhood maltreatment (CM) and substance use disorders (SUD), as also described in Capusan et al., 2021 and Perini et al., 2023. However, there were no group differences across the measures described below, so all analyses were collapsed across groups for a total N = 101 (see Table 1 for demographic information).

Participants were initially recruited based on presence or absence of prospective documental childhood maltreatment (CM) or substance use disorder (SUD) histories. The first group had both CM and lifetime SUD (CM+SUD, n = 28); the second group had CM without lifetime SUD (CM only, n = 24); the third group consisted of a healthy control group with neither CM nor lifetime SUD (control, n = 24); finally, the fourth group consisted of a clinical control group with lifetime SUD but no documented CM (SUD only, n = 25). All CM exposed participants (CM only and CM+SUD) consisted of former patients in a specialized treatment unit for children and adolescents exposed to physical and/ or sexual abuse and/ or severe neglect referred by the child protective services[1]. The Swedish personal identification number allowed the identification and long-term follow up of these former CM treatment unit patients, now young adults, using the regional health care register for Östergötland County, Sweden (n = 470). Sixty-five former CM treatment unit patients with both documented CM exposure and documented contact with SUD clinics were eligible. For each of these participants, we identified sex/age matched CM exposed eligible individuals with no lifetime SUD (n = 140), and sex/age matched individuals with lifetime SUD but with no recorded CM exposure (n = 106). Controls with lifetime SUD but no documented CM were recruited using the regional health care register and through advertisements from addiction clinics in the Region of Östergötland. Sex and age matched healthy controls with no documented SUD or CM were recruited through advertising among students at Linköping University and social media. Participants completed breath and urine screens for alcohol and drug use prior to laboratory sessions.

During the screening session, participants underwent a psychiatric clinical assessment by a trained research nurse or study physician, and a structured Mini International Neuropsychiatric Interview (MINI-7) [2] for DSM-5, Swedish version. Lifetime SUD was identified using the regional health care register and contact with addiction clinics, and current SUD was assessed using the MINI, self-reported current problems and urine drug screens. Controls were at this point excluded if CM was identified in their medical records, during clinical assessment or with the MINI-7 post-traumatic stress disorder module. After determining eligibility, participants received research information, provided written informed consent, and completed self-report questionnaires. Questionnaires assessed self-reported CM, using the Childhood Trauma Questionnaire-Short Form (CTQ)[3,4]. Current alcohol and drug use severity were assessed using the Alcohol Use Disorders Identification Test (AUDIT)[5] and the Drug Use Disorders Identification Test (DUDIT)[6]. Emotion regulation and personality traits were assessed using the Difficulties in Emotion Regulation Scale (DERS-16)[7].

#### Human Fear Conditioning

Upon arrival at the lab, participants were fitted with an intravenous catheter for blood sample collection for subsequent analysis of endocannabinoid levels and prepared for psychophysiological recordings via application of facial electromyography (EMG) recording electrodes. Two 4mm silver/silver chloride bipolar recording electrodes placed over the orbicularis oculi muscle (“orbicularis”, below the eye) on the left side of the face and an 8mm ground electrode placed on the forehead near the hairline. Sites were cleaned with alcohol and lightly abraded and electrodes were reapplied on any site with impedance over 20kΩ (measured with a Model 1089 MK III Checktrode; UFI, Morro Bay, CA, USA). EMG data collection and fear conditioning took place as previously described [8,9]. EMG signals were amplified, filtered through a 10-500 Hz band pass and 50 Hz comb band stop filter, digitized at 1 kHz, re-filtered, rectified, and integrated over 20ms using EMG100C amplifiers, MP150 Data Acquisition system and Acqknowledge software from Biopac Systems (Biopac Systems, Inc, Camino Goleta, CA, USA). The fear conditioning task consisted of Habituation, Acquisition, and Extinction, with the first 4 trials of Extinction serving as the CS discrimination and Recall test [10]. Auditory stimuli were presented via Sennheiser Model HD-25 headphones. Tasks were presented using Presentation Software (Neurobehavioral Systems; Berkley, CA, USA). Habituation took place first and consisted of presentations of all conditioned stimuli and contexts, but no unconditioned stimulus was delivered. In Acquisition, subjects were exposed to 8 presentations of the conditioned stimulus (lamp color, CS+) predicting an unpleasant sound (nails across the chalkboard; US) [11] and 8 presentations of another CS (different lamp color, CS-) which was not followed by any aversive stimulus. After a 10min break, Recall took place in a different context consisting of a digital photograph of a different room than that used during acquisition (reception waiting room or an office). Subjects were exposed to the same CS+ and CS- but no US. Here, we assessed CS discrimination at the start of early Extinction by presenting subjects with 2 CS- and 2 CS+. A startle probe (50ms burst of white noise) was used to elicit eye-blinks during each CS and inter-trial intervals, measured as the peak-to-peak orbicularis occuli EMG value in the 21-150ms window after probe onset [12]. To account for individual differences in startle magnitude [12], startle responses to the CS+ were standardized to the mean startle response during ITI/rest trials, calculated as follows: [average startle to CS+]/[average startle during rest], with values > 1 indicating potentiation of the startle response [8,9,13]. All startle responses were visually inspected by trained, blinded raters and scored as missing values if a voluntary blink occurred just before, during, or after probe onset, or if there were any other artifacts obscuring the response [12].

**Endocannabinoid analysis**

An indwelling catheter was used to collect blood samples for analysis of baseline endocannabinoids, calculated as the average of two timepoints. The eCBs anandamide (AEA) and 2-arachidonoylglycerol (2-AG) were extracted and analyzed using liquid chromatography tandem mass spectrometry (LC-MS/MS), as previously published [8]. Briefly, 300 µL of serum was thawed and vortexed, and 30 µL of a mixture containing the deuterated internal standard (AEA-d4, 50 nM; 2AG-d5, 1000 nM; Cayman Chemicals, Ann Arbor, MI, USA) was added to each serum sample. C8 Octyl SPE columns (6 mL, 200 mg; Biotage; Uppsala, Sweden) were used for lipid extraction. Prior to transferring the samples, the C8 Octyl SPE columns were activated with 1ml Methanol (Merck, Darmstadt, Germany) and washed with 1mL MilliQ-H2O using a Biotage ® Pressure+ 48 machine. After the samples were added, the columns were washed with ACN (20% with 0.1% TFA) and samples were eluted with ACN (80% with 0.1% TFA). The eluates were evaporated to dryness in a SpeedVacc (Thermo Fisher, Ann Arbor, MI, USA) and stored in -80 °C until analysis. On the day of the analysis, the samples were reconstituted in 30µl Mobile phase A (methanol-milliQ water-acetonitrile (4/4/2) (v/v/v) with 0.1 % (v/v) formic acid and 1g/L ammonium acetate), then vortexed and transferred into glass vials designed for the LC-MS/MS. The injection volume was 10 µL. We used an LC-MS/MS system consisting of a Thermo Scientific Accela AS auto sampler and Accela 1250 pump coupled to a Thermo Scientific TSQ Quantum Access max triple quadrupole mass spectrometer with a HESI II probe as ionization source. LC was performed using gradient elution with mobile phase A, and mobile phase B (containing methanol-ACN (7/3) (v/v) with 0.1% (v/v) formic acid and 1g/L ammonium acetate). The gradient elution was applied with a constant flow of 250 µL/min. We started with 100% mobile phase A during the first 1.5 min and followed this using a linear increase towards 100% mobile phase B, which was achieved after 9 min in total. Between the 11th and 12th min the gradient changed linear to 100% mobile phase A, which was maintained for 1 min. An Xbridge C8 analytical column (2.1 mm × 150 mm) with the particle size 2.5 µm obtained from Waters (Dublin, Ireland) was used. We used the following selected reaction monitoring (SRM) (m/z) transitions: 348.3/ 62.4 and 379.3/287.3 for AEA and 2-AG, respectively. For the corresponding internal standards, we used the following transitions: 352.3/ 62.4 and 384.3/287.3 for AEA-d4 and 2-AG-d5, respectively. The linearity of the measuring ranges was assessed with standard curves ranging from 1-25 nM for AEA and and 50-1250 nM for 2-AG in duplicate. The linearity of the standard curves was R2 ≥ 0.96 for all analytes. Isotopic dilution was used for quantification of the analytes, performed according to their area ratio of their corresponding deuterated internal standard signal area. 2-AG was quantified as the sum of 1- and 2-AG, which has been reported by others [14]. Linear regression and $X^2$ weighting were applied. Undetected levels were considered as 0 nM. Xcalibur® (version 2.1, Thermo Scientific) software was used for peak integration and quantification. Endocannabinoid values were log transformed due to non-normality of the distribution; these transformed values were used in all subsequent analyses. Baseline differences in endocannabinoids were added as co-variates in analysis of brain, behavioral, and self-report measures.

**Magnetic Resonance Imaging**

Anatomical and functional blood oxygen-level-dependent (BOLD) data were collected on a Siemens MAGNETOM Prisma 3T MRI scanner (Siemens healthcare AB, Stockholm, Sweden) equipped with a 64-channel head coil. Resting state images were collected using an echoplanar imaging (EPI) sequence with the following parameters: TR=901 ms; TE=30 ms; voxel size=3.0 mm isotropic; no slice gap; FOV=192 mm x 192 mm x 144 mm; flip angle=63°, number of volumes=800. Anatomical data was collected via T1-weighted imaging sequence: TR=2300 ms; TE=2.36 ms; voxel resolution=0,9 mm isotropic; no slice gap; flip angle=8°; FOV=263 mm x 350 mm x 350 mm. EPI images were de-spiked, slice-time corrected, smoothed (4 mm) and motion-corrected.

Preprocessing and statistical analysis of resting state data were performed with the Analysis of Functional Neuro Images (AFNI) software v18.3.16 [15]. A Freesurfer-based parcellation was performed on T1-weighted data using the function recon-all and used for tissue-based regression to allow for modelling of non-BOLD signal fluctuations. BOLD images were de-spiked, slice-time and motion-corrected, spatially transformed to MNI template space and smoothed (4 mm). Volumes exceeding 2 mm motion censoring and 0.05 outlier fraction were not included in the time-series regression. Preprocessed BOLD time-series data were then entered in a regression using the 3dDeconvolve function and head motion effects were accounted for by adding the motion parameters and their derivatives as regressors of no interest. Covarying signals between gray and white matter regions were removed from the residuals using the data driven APPLECOR method [16].

Seed-based connectivity analyses were performed by entering seed time course as predictor in regression analyses. Right and left amygdala masks were used as independent seeds. To guarantee that voxels in the masks were not affected by signal loss, amygdala masks were multiplied on individuals' EPI-masks surviving censoring. The target region was defined by using the Matthew Glasser atlas for the dlPFC (bilateral areas 46, 9-46d, anterior and posterior 9-46v, and anterior 9; Figure S5). For each participant (N = 88), amygdala time course was entered as predictor in a regression analysis on preprocessed data, using 3dDeconvolve. Resulting 3D volumes with beta coefficients for right/left amygdala were assessed for covariance with peripheral measures of endocannabinoid function and behavioral measures of fear conditioning using the AFNI function 3dttest++ and with 2AG and CS discrimination (CS+ vs CS-) at early Extinction as covariates in two separate analyses. To validate the results, beta coefficients from the significant cluster were then extracted for each participant and entered in regression analyses with the corresponding covariate as predictor. Regression analyses were performed using SPSS version 25.

**Rodent Surgeries**

As previously described on [17], mice were initially anesthetized in an induction chamber with 5% isoflurane and then transferred to the stereotax (Kopf Instruments, Tujunga, CA) and kept under 2% isoflurane anesthesia through the duration of the surgery. Skull hairs were trimmed and the exposed skin was prepped with alternating alcohol and iodine scrub. The skull was exposed via a midline incision. The incision site was treated with the local anesthetic benzocaine (Medline Industries, Brentwood, TN). For all surgeries, we used digital software (NeuroStar; Tübingen, Germany) to control movement of the stereotaxic frame arm. A 10 mL microinjection syringe (Hamilton Co., Reno, NV) driven by a Micropump Controller (World Precision Instruments, Sarasota, FL) attached to the stereotaxic frame was used to inject AAVs encoding GCaMP7f under control of the synapsin promoter (AAVrg-syn-jGCaMP7f-WPRE with titer of 1.8 × 10^13 vg/mL; Addgene, Watertown, MA) into PL at 2 dorsoventral levels (AP:+2.42 mm, ML: + 0.35 mm, DV: 1.6 mm and 1.9 mm), 300 nL per level. After allowing for diffusion of the virus, the needle was withdrawn, and a 0.5 mm diameter needle attached to a vacuum line was lowered stereotaxically to a depth of 1.55 mm above PL (AP:+2.42 mm, ML: + 0.35 mm, DV: 1.6 mm and 1.55 mm) to create a tract. Thereafter, a 0.5 mm diameter GRIN lens (Inscopix, Palo Alto, CA) was slowly lowered stereotaxically into PL at a depth of 1.8 mm DV. The GRIN lens was affixed to the exposed skull using Metabond (Parkel, Edgewood, NY). Following completion of the surgery, 10 mg/kg ketoprofen (AlliVet, St. Hialeah, FL) was administered as an analgesic, and again as post-operative treatment 24 and 48 h after the surgery. In some cases, animals were allowed to recover for at least 2 weeks, after which a baseplate used to dock the miniaturized microscope was installed over the lens at an empirically optimized working distance. In other cases, no baseplate was installed as the GRIN lenses already had integrated baseplates.

**Rodent Behavior and recordings**

Auditory fear conditioning and recall testing were performed in the same operant chambers (Coulbourn Instruments, Allentown, PA) located in sound-attenuating cubicles (Coulbourn Instruments, Allentown, PA) throughout all phases of the experiment. The floor of the chambers consisted of a grid of parallel stainless-steel bars that allow delivery of a scrambled electric foot shock. Between experiments the grid, chamber wall, and pan were cleaned with 70% ethanol and dried thoroughly. On Day 1, mice were conditioned to tones in Context A (CtxA) using various variants of fear conditioning as depicted in Figure 2. In all cases of auditory fear conditioning, all mice were exposed to 1 habituation tone (no foot shock) and 8 conditioned tones (CS+) with 15 s duration co-terminating with a 1 s duration 0.4 mA electric foot shock. CS+ consisted of either 2kHz tones or white noise which were counterbalanced across experiments to control for stimulus effects. For the differential fear conditioning and partial differential conditioning experiments, mice were exposed to tones of opposite identity to that of the CS+ and which were not paired with foot shocks, denoted CS-. On Day 2, mice were injected intraperitonially with DO34 (Glixx Laboratories, Hopkinton, MA) or vehicle at a dose of 50 mg/kg in a formulation containing ethanol:Kolliphor:saline (1:1:18; Sigma-Aldrich, St. Louis, MO). 2 hours later, mice were returned to Context B (CtxB) which consisted of the same conditioning chambers modified by covering the floor grid with white vinyl covered plexiglass and inserting a transparent plexiglass insert with alternating white vinyl stripes to form a semi-cylindrical chamber. A container containing 10% vanilla extract (McCormick, Cockeysville, MD) was left inside the cubicles to provide a distinct odor from that in CtxA. Mice were then presented with CS- and CS+ tones. In the case of mice not exposed to CS- tones on Day 1, mice were presented with Novel Tones (NT; consisting of the opposite identity of the CS+) and CS+ as depicted in Figure 2.

For contextual fear conditioning experiments, mice were exposed to 2 sessions in CtxA, 270 s in duration, 1 per day, with 2 0.4 mA foot shocks, 2 s in duration. On Day3, mice were injected with either DO34 or vehicle and exposed to either CtxA or novel context (NCtx) for 240 seconds in a 2x2 design. Data was analyzed for seconds 120 to 240 of context exposure. NCtx consisted of the same arrangement used for CtxB in the experiments previously described.

For our neuronal calcium activity recordings, mice were habituated to the miniaturized epifluorescence microscope (Inscopix, Palo Alto, CA) for at least 2 days. On Day 1, the microscope was attached to the mice and were fear conditioned as depicted in the diagram in Figure 3. On Day 2, the microscope was attached to the mice and baseline calcium activity was recorded for 15 minutes with mice remaining in their home cage after which the microscope was detached, and mice were injected with either vehicle or DO34. After 2 hours, the microscope was reattached, and post-injection activity was recorded for 15 minutes after which mice were then transferred to CtxB and exposed to NT and CS+ presentations. Upon completion of the behavioral session, mice were disconnected from the microscope and returned to their home cage. As previously described, calcium activity data was acquired at a frame rate of 10 Hz [17]. LED power, gain, and lens focus were empirically adjusted to maximize the quality of the recordings while minimizing the LED power used. Data was acquired continuously throughout the duration of the session.

All the fear conditioning experiments were controlled using FreezeFrame (Actimetrics, Wilmette, IL). For calcium activity recordings, the calcium imaging data acquisition system (Inscopix, Palo Alto, CA) was interfaced with the experimental control software using custom made RJ12 connector to BNC connector cables to deliver TTL pulses synchronizing the start of the behavioral session with the start of data acquisition and flagging tone deliveries within the recordings.

**Rodent Data Analysis**

Upon completion of the experiments, freezing data was obtained from FreezeFrame. The amount of time spent freezing to the tones or context was expressed as a percentage of the total interval duration. Tone trials are presented in blocks of 2.

Calcium imaging data was analyzed as previously described [17,18]. Briefly, recordings were spatially down sampled by a factor of 2. Resulting videos were then bandpass filtered and motion corrected using Inscopix Data Processing software V1.3. Subsequently, individual neurons were identified, and their calcium traces extracted using constrained non-negative matrix factorization for endoscopy (CNMFe)[19] algorithm. We used the resulting non-deconvolved traces for our analyses. Traces and corresponding identified neurons were individually verified for quality and traces of falsely identified neurons were excluded from further analysis. The resulting traces were aligned to the tone onset and traces were binned into 1 s intervals. 2 consecutive tone trials were averaged together into blocks of 2 for our analyses. To assess for tone induced activity changes, the data was normalized by applying the Z-transform using the 15 s pre-tone interval as the normalization baseline. Traces with a Z-score >3 ($p = 0.01$, two-tailed) for any 2 consecutive bins following tone onset during the tone period were considered (+) Responsive and Z-score <-3 ($p = 0.01$, two-tailed) were considered (-) Responsive. Longitudinal registration across 2 days or between pre-post drug injection baseline sessions was conducted using Inscopix Data Processing software using 0.5 minimum correlation as threshold. The resulting aligned traces were individually reviewed for continuity across the 2 sessions and analyzed as described above.

Agglomerative hierarchical clustering was conducted using the MATLAB clustergram function. The Z-scored traces for the NT and CS+ were concatenated together with each row corresponding to the traces from a single neuron. Linkage was then calculated using Ward's method (inner squared distance) with Euclidean distance for the metric. The algorithm pairs neurons in close linkage proximity and then clusters the resulting clusters with others in close proximity, this process is repeated iteratively until a hierarchical tree (a dendrogram) is built. 3% of the maximum linkage was used as a threshold to determine the different clusters which are identified by distinct colors within the dendrogram and above the associated heatmap.

Calcium spike data was also used to obtain baseline and post-drug spontaneous activity rates. For this, we took the total amount of spikes in the session and divided that number by the total session duration (15 minutes) in seconds. The spike data was obtained from the deconvolved traces using the Oasis package from CNMFe.

1. Capusan AJ, Gustafsson PA, Kuja-Halkola R, Igelstrom K, Mayo LM, Heilig M. Re-examining the link between childhood maltreatment and substance use disorder: a prospective, genetically informative study. *Mol Psychiatry*. Jul 2021;26(7):3201-3209. doi:10.1038/s41380-021-01071-8
2. Sheehan DV, Lecrubier Y, Sheehan KH, et al. The Mini-International Neuropsychiatric Interview (M.I.N.I.): the development and validation of a structured diagnostic psychiatric interview for DSM-IV and ICD-10. *J Clin Psychiatry*. 1998;59 Suppl 20:22-33;quiz 34-57.
3. Bernstein DP, Stein JA, Newcomb MD, et al. Development and validation of a brief screening version of the Childhood Trauma Questionnaire. *Child Abuse Negl*. Feb 2003;27(2):169-90. doi:10.1016/s0145-2134(02)00541-0

4. Bernstein DP, Fink L, Handelsman L, et al. Initial reliability and validity of a new retrospective measure of child abuse and neglect. *Am J Psychiatry*. Aug 1994;151(8):1132-6. doi:10.1176/ajp.151.8.1132
5. Babor TF, Higgins-Biddle JC, Saunders JB, Monteiro MG, Organization WH. *AUDIT: the alcohol use disorders identification test: guidelines for use in primary health care*. 2001.
6. Berman AH, Bergman H, Palmstierna T, Schlyter F. Evaluation of the Drug Use Disorders Identification Test (DUDIT) in criminal justice and detoxification settings and in a Swedish population sample. *Eur Addict Res*. 2005;11(1):22-31. doi:10.1159/000081413
7. Bjureberg J, Ljotsson B, Tull MT, et al. Development and Validation of a Brief Version of the Difficulties in Emotion Regulation Scale: The DERS-16. *J Psychopathol Behav Assess*. Jun 2016;38(2):284-296. doi:10.1007/s10862-015-9514-x
8. Mayo LM, Asratian A, Linde J, et al. Protective effects of elevated anandamide on stress and fear-related behaviors: translational evidence from humans and mice. *Mol Psychiatry*. May 2020;25(5):993-1005. doi:10.1038/s41380-018-0215-1
9. Mayo LM, Asratian A, Linde J, et al. Elevated Anandamide, Enhanced Recall of Fear Extinction, and Attenuated Stress Responses Following Inhibition of Fatty Acid Amide Hydrolase: A Randomized, Controlled Experimental Medicine Trial. *Biol Psychiatry*. Mar 15 2020;87(6):538-547. doi:10.1016/j.biopsych.2019.07.034
10. Milad MR, Orr SP, Pitman RK, Rauch SL. Context modulation of memory for fear extinction in humans. *Psychophysiology*. Jul 2005;42(4):456-64. doi:10.1111/j.1469-8986.2005.00302.x
11. Neumann DL, Waters AM, Westbury HR, Henry J. The use of an unpleasant sound unconditional stimulus in an aversive conditioning procedure with 8- to 11-year-old children. *Biol Psychol*. Dec 2008;79(3):337-42. doi:10.1016/j.biopsycho.2008.08.005
12. Blumenthal TD, Cuthbert BN, Filion DL, Hackley S, Lipp OV, van Boxtel A. Committee report: Guidelines for human startle eyeblink electromyographic studies. *Psychophysiology*. Jan 2005;42(1):1-15. doi:10.1111/j.1469-8986.2005.00271.x
13. Walker DL, Davis M. Quantifying fear potentiated startle using absolute versus proportional increase scoring methods: implications for the neurocircuitry of fear and anxiety. *Psychopharmacology (Berl)*. Nov 2002;164(3):318-28. doi:10.1007/s00213-002-1213-0
14. Balvers MG, Verhoeckx KC, Witkamp RF. Development and validation of a quantitative method for the determination of 12 endocannabinoids and related compounds in human plasma using liquid chromatography-tandem mass spectrometry. *J Chromatogr B Analyt Technol Biomed Life Sci*. May 15 2009;877(14-15):1583-90. doi:10.1016/j.jchromb.2009.04.010
15. Cox RW. AFNI: software for analysis and visualization of functional magnetic resonance neuroimages. *Comput Biomed Res*. Jun 1996;29(3):162-73. doi:10.1006/cbmr.1996.0014
16. Marx M, Pauly KB, Chang C. A novel approach for global noise reduction in resting-state fMRI: APPLECOR. *Neuroimage*. Jan 1 2013;64:19-31. doi:10.1016/j.neuroimage.2012.09.040
17. Patel S, Johnson K, Adank D, Rosas-Vidal LE. Longitudinal monitoring of prefrontal cortical ensemble dynamics reveals new insights into stress habituation. *Neurobiol Stress*. Sep 2022;20:100481. doi:10.1016/j.ynstr.2022.100481
18. Marcus DJ, Bedse G, Gaulden AD, et al. Endocannabinoid Signaling Collapse Mediates Stress-Induced Amygdalo-Cortical Strengthening. *Neuron*. Mar 18 2020;105(6):1062-1076 e6. doi:10.1016/j.neuron.2019.12.024
19. Zhou P, Resendez SL, Rodriguez-Romaguera J, et al. Efficient and accurate extraction of in vivo calcium signals from microendoscopic video data. *Elife*. Feb 22 2018;7doi:10.7554/eLife.28728
